# Measurement and Control of Crossed Potentials in a Flavoprotein

**DOI:** 10.1101/2025.09.13.676020

**Authors:** Benjamin J. Jones, Sarah Elhajj, Brett Haynes, Sichu Wang, Ian O’Connor, Samer Gozem, Brandon L. Greene

**Affiliations:** Department of Chemistry and Biochemistry, University of California Santa Barbara, Santa Barbara, CA USA 93106; Department of Chemistry, Georgia State University, Atlanta, GA USA 30302; Interdisciplinary Program in Quantitative Biosciences, University of California Santa Barbara, Santa Barbara, CA USA 93106

## Abstract

Flavoproteins are versatile redox-active biomolecules enabling a multitude of metabolic processes. Their versatility stems from the tunability of the flavin cofactor’s one- and two-electron reduction potentials via interactions with the protein scaffold, which have dramatic influence on reactivity. Although several mechanisms have been proposed to explain how the flavin-binding pocket modulates redox thermodynamics, few have been validated through quantitative experiments. In this study, we investigate how the flavin N5 environment influences the redox properties of the flavin mononucleotide cofactor in the “improved” light-oxygen-voltage (iLOV) sensing protein using site-directed mutagenesis, redox titrations, and hybrid quantum mechanical molecular mechanical (QM/MM) methods combined with classical alchemical free energy simulations. Mutating the residue Q_103_, which interacts with the flavin N5 and O4′ atoms in the X-ray crystallographic structure, exerts a modest < 35 mV effect on the overall two-electron reduction potential, but significantly alters the potential separation of the two one-electron couples (potential crossing) by up to 168 mV. QM/MM and free energy calculations reveal that water penetration into the flavin binding pocket near N5 and O4′ largely explains the trend in reduction potentials among the mutants. The results suggest a molecular mechanism of flavin tuning in which hydrogen bonding to the neutral semiquinone, either directly by the side-chain or a protein-penetrating water, contributes significantly to the potential crossing. These findings establish quantitative experimental benchmarks for theoretical models and advance a molecular mechanism for redox tuning in flavoproteins.

## INTRODUCTION

Flavins play crucial roles in biology as redox-active enzyme cofactors and electron transfer agents.^1–3^ The versatility of the flavin cofactor arises from its ability to mediate both one- and two-electron transfer reactions.^4,5^ The oxidized (OX) cofactor can be reduced by one electron to generate a radical semiquinone (SQ) state or accept two electrons to form the fully reduced hydroquinone (HQ) state (**Figure 1A**). These redox transformations can be proton-coupled, resulting in either neutral (N) or anionic (A) forms of the SQ and HQ states. The reduction potentials of these couples (E°_OX/SQ_ and E°_SQ/HQ_) are highly sensitive to the local protein environment, allowing enzymes to tune flavin redox chemistry for optimal thermodynamic efficiency and kinetic performance.^6^

**Figure 1.**
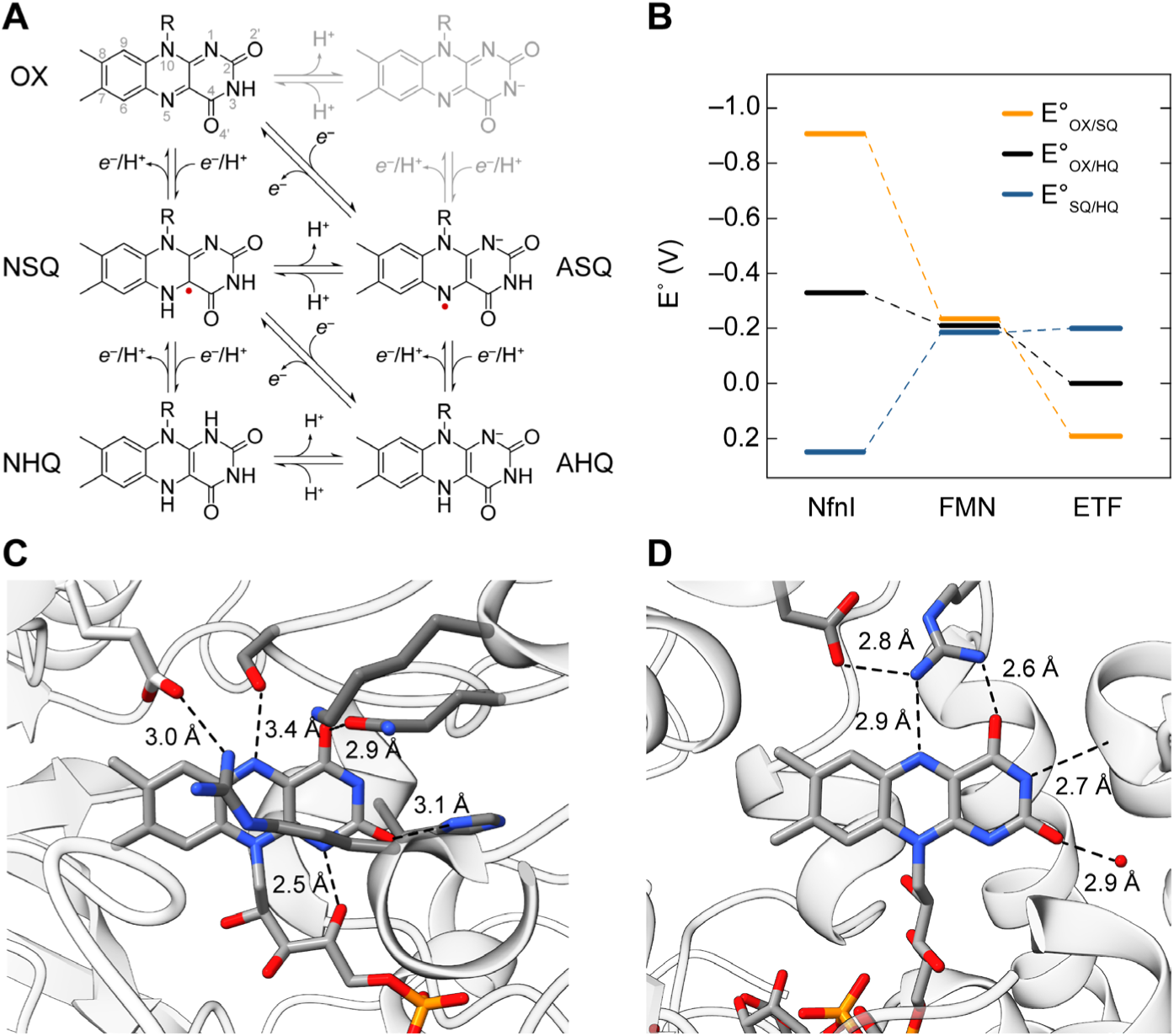
Flavin redox chemistry and tuning in flavoproteins. **A** Redox and protonation states of the flavin isoalloxazine ring. **B** Reduction potentials for the three redox couples of *Pyrococcus furiosus* (*Pf*) NfnI, free FMN, and *Methylophilus methylotrophus* (*Mm*) ETF, demonstrating the span of modulation the protein environment exerts on flavin redox chemistry. **C** Interactions of the *Mm* ETF (PDB ID: 1O96) FAD cofactor with the protein environment. **D** Interactions of the *Pf* NfnI (PDB ID: 5JFC) FAD cofactor with the protein environment. Both proteins have a D/E→R salt bridge where a R guanidinium nitrogen is within 4.5 Å of the flavin N5. The N3 of *Pf* NfnI is hydrogen bonded to a valine backbone carbonyl and N1 is hydrogen bonded to the substrate nicotinamide amide oxygen of NADPH (3.2 Å, not shown).

In solution at neutral pH, free flavin mononucleotide (FMN) exhibits “crossed” potentials: the reduction potential of the OX/NSQ couple (E°′_OX/NSQ_ = ‒238 mV vs. NHE) is more negative than the subsequent NSQ/AHQ couple (E°′_NSQ/AHQ_ = ‒172 mV vs. NHE). As a result, reduction occurs as concerted two-electron process, and the NSQ is unstable (**Figure 1B**).^7^ Flavoproteins can stabilize or destabilize an SQ intermediate to achieve a wide range of redox behaviors. For example, consider flavin-based electron bifurcating enzymes, including the NADPH-dependent ferredoxin:NAD^+^ oxidoreductases (Nfn), heterodisulfide reductases (Hdr), and bifurcating electron transfer flavoproteins (Etf), which couple the oxidation of a moderate two-electron donor to the one-electron reduction of both an exergonic and endergonic acceptor.^8,9^ In these enzymes, electron bifurcation occurs at a flavin adenine dinucleotide (FAD) cofactor with significantly crossed one-electron potentials and an unstable (A)SQ intermediate, a feature proposed to be critical for electron bifurcation.^10,11^ In the *Pyrococcus furiosus* NfnI, which catalyzes the NADPH-dependent reduction of both NAD^+^ (exergonic) and ferredoxin (endergonic), the potential splitting (ΔE = E°_OX/SQ_ ‒ E°_SQ/HQ_) between the E°_OX/ASQ_ and E°_ASQ/AHQ_ couples is estimated to be ‒1.2 V.^12^ By contrast, the *Methylophilus methylotrophus* non-bifurcating electron transfer flavoprotein (ETF), which facilitates electron transfer from trimethylamine dehydrogenase to the electron transport chain,^13^ exhibits normally ordered potentials; the E°_OX/ASQ_ midpoint potential is +196 mV and a much lower ‒197 mV for the E°_ASQ/AHQ_, and a corresponding ΔE of +393 mV. The higher potential OX/ASQ couple is well matched to the electron donor, an iron-sulfur cluster of trimethylamine dehydrogenase with a reduction potential of +79 mV, thus enabling facile one-electron transfer, and the E°_ASQ/AHQ_ couple is metabolically irrelevant.^14^

Comparison of the redox behavior of *P. furiosus* NfnI and *M. methylotrophus* ETF reveals that the flavin environment can tune the two one-electron potentials by more than 1 V (> 100 kJ/mol), ranging from strongly crossed to normally ordered. Yet a simple inspection of their respective flavin protein environments does not immediately reveal the origin(s) of these differences (**Figure 1C** and **1D**). Hence, how the flavin environment achieves this control remains poorly understood. It is likely that the combined effects of sterics,^15,16^ hydrogen bonding,^17,18^ aromaticity,^17,19^ and coulombic effects^20–22^ contribute to this tuning, and are known to modulate flavin redox properties in flavoproteins. Yet, definitive experimental evidence isolating scaffold contributions to flavin redox tuning remains elusive. Such structure-function relationships have been experimentally challenging to demonstrate due to the weak binding of flavin cofactors, variations that can be seen by post-translational modifications or substrate/product/effector binding, and the difficulty of measuring the effects on crossed potential flavins, where the SQ intermediate is unstable.

Here, we leverage the “improved” light-oxygen-voltage sensing (iLOV) protein as a model flavoprotein to begin to dissect the role of the protein scaffold in controlling redox behavior of a crossed potential flavin.^23,24^ Using site-directed mutagenesis, we altered residue Q_103_ (residue 489 in the original *Arabidopsis thaliana* phototropin 2 numbering scheme), which forms a hydrogen bond with the flavin N5/O4′ in the oxidized state.^24^ Potentiometric titrations, spectrophotometric comproportionation, and electron paramagnetic resonance (EPR) spectroscopy demonstrate that Q_103_ variants primarily shift E°_OX/NSQ_, while minimally affecting E°_NSQ/NHQ_, thereby altering ΔE. Hybrid quantum mechanical molecular mechanical (QM/MM) and classical alchemical free energy simulations reproduce these experimental trends and provide a mechanistic basis; hydrogen bonding to N5/O4′ by penetrating water molecules from the solvent stabilizes the NSQ state, but not the OX state, increases E°_OX/NSQ_. Taken together, these results establish that the protein scaffold is capable of independent tuning of flavin one-electron chemistry by modulating N5 protonation and N5/O4′ hydrogen bonding, offering new strategies for rational design and mechanistic interpretation of flavoprotein redox behavior.

## EXPERIMENTAL DETAILS

### Materials

The iLOV gene within a pQE80L vector backbone was obtained from the Mukherjee lab and cloned into a pSUMO vector as reported previously.^25,26^ The 6× polyhistidine-tagged ubiquitin-like protease 1 (ULP1) was expressed and purified as reported previously.^27,28^ Primers for PCR were ordered from Integrated DNA Technologies (IDT). DpnI, Phusion HF polymerase, chemically competent DH5-α and BL21 (DE3) *Escherichia coli*, the Qiagen QIAprep Spin Miniprep and Gel Extraction kits, and the NEBuilder HiFi DNA Assembly kit were purchased from New England Biolabs (NEB). IPTG was purchased from Merck. Sephadex G-25 fine resin, LB Miller broth, NaH_2_PO_4_ (P_i_), dithiothreitol (DTT), phenylmethylsulfonyl fluoride (PMSF), bovine xanthine oxidase (XO), xanthine, 4-hydroxy-(2,2,6,6-tetramethylpiperidin-1-yl)oxyl (4-hydroxy-TEMPO), sodium hydrosulfite (dithionite, NaDT), and Amicon Ultra centrifugal filters were purchased from Millipore Sigma. FMN sodium salt, HisPur nickel nitrilotriacetic acid (Ni-NTA) resin, bicine, safranine O, methyl viologen (MV^2+^), 96-well plates, Pierce 660 nm protein assay kit, amber NMR tubes, and Imperial protein stain were obtained from Thermo Fisher Scientific. Bradford reagent was purchased from VWR. Kanamycin sulfate was obtained from Genesee Scientific. Riboflavin was obtained from Acros Organics. Microseal ‘B’ 96-well plate seals were obtained from BioRad. The single-color, cold, mounted, light-emitting diode (LED) with a wavelength of 450 nm and a bandwidth of 18 nm, KG-1 and KG-2 glass filters were obtained from ThorLabs. The pyrolytic graphite edge-plane (PGE) electrode, platinum wire counter electrode, Ag/AgCl (3M) reference electrode, and saturated calomel reference electrode (SCE) were obtained from Bioanalytical Systems (BASi) research products. The SP-200 single channel potentiostat and FC-45 Faraday cage were obtained from BioLogic. Alumina polishing pads and 0.05, 0.3, 1.0 μm sized alumina were obtained from Buehler. Precision quartz (0.40 mm wall) EPR tubes were obtained from SP Wilmad. Nanopure water was obtained via a Barnstead E-Pure ultrapure water system (>17 MΩ/cm) and was used for all solutions.

### Methods

Construction of the pSUMO-iLOV plasmid vector, growth and aerobic purification of (H)_6_-SUMO-iLOV and mutants, ULP1 digestion, purification of iLOV, and loading and quantitation of FMN were conducted as described previously.^26^ Total protein yields and associated FMN content is provided in **Supplemental Information Table S1** and protein purity gels are shown in **Supplemental Information Figure S1**.

### Site-Directed Mutagenesis of iLOV

Mutagenic primers used for site-directed mutagenesis to generate Q_103_X (X = G, A, R, K, and D) iLOV are reported in **Supplemental Information Table S2**, and mutagenesis was carried out via the polymerase chain reaction (PCR) per the NEB protocol. The PCR products were transformed into *E. coli* DH5α and the cells were plated onto LB/agar plates supplemented with 50 mg/L kanamycin and grown overnight. The following day, colonies were picked for growth in small 5 mL cultures. Mutant plasmids were purified by DNA miniprep and sent for nanopore sequencing at UC Berkeley’s DNA sequencing facility. Upon confirmation of the correct sequences, mutant plasmids were used to transform *E. coli* BL21(DE3) cells for protein expression and growth as described previously.^26^

### Quantum Yields, Excitation, and Emission Spectra

Samples of iLOV were prepared in 50 mM P_i_ 200 mM NaCl at pH 7.50 to an absorbance of 0.05 at 450 nm to minimize inner filter effects, and verified on an Agilent Cary 60 UV-vis spectrophotometer in triplicate. The emission spectra of each variant between 470 and 900 nm was collected in a Duetta Spectrometer (Horiba Scientific) using an excitation wavelength of 450 nm and samples were thermostated at 25 °C. An excitation bandpass was set to 5 nm, while the emission bandpass was set to 1 nm with an emission increment of 0.5 nm. A total of 50 detector accumulations each with an integration time of 300 ms were averaged to obtain final emission spectra. The quantum yield of the sample was calculated using **eq. 1**.

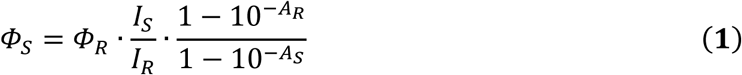

Here, *Φ_R_* is the quantum yield of the sample unknown (iLOV and Q_103_X mutants), *Φ_R_* is the quantum yield of some reference, (in this case riboflavin with a reported quantum yield of 0.26),^29^ *I_S_* and *I_R_* represent the integrated emission intensity of the sample and reference, respectively, and *A_S_* and *A_R_* represent the absorbance of the sample and reference, respectively. The standard deviations of the *Φ_S_* were propagated by both the standard deviation of the absorbance measurement and the integrated emission intensity.

Excitation spectra were collected on the same instrument scanning excitation wavelengths from 300 to 500 nm with an excitation step of 1 nm. The emission wavelength was fixed at 520 nm. Both the excitation and emission bandpass were set to 5 nm with an integration time of 50 ms. A total of 10 detector accumulations were used for each data point.

### Photoluminescence Lifetimes

Samples of iLOV and Q_103_X mutants were diluted to an absorbance of 0.05 at 400 nm to minimize inner filter effects. Fluorescence lifetime measurements were performed using time-correlated single photon counting (TCSPC). An approximately 100 fs excitation pulse with wavelength of 400 nm was generated by doubling the fundamental frequency of Ti:Sapphire laser (Spectraphysics Tsunami) pulses in a commercial optical harmonics generator (Inrad/Coherent). The laser repetition rate was reduced by a home-made acousto-optical pulse picker in order to avoid saturation of the chromophore. The TCSPC detection system was equipped with a single-photon counting avalanche photodiode module (Photon Micro Devices) and electronics board (PicoQuant PicoHarp 300) with an instrument response time < 50 ps. A triggering signal for the TCSPC board was generated by sending a small fraction of the excitation beam onto a fast (400 MHz bandwidth) Si photodiode (Thorlabs Inc.). The fluorescence signal was collected in a 90° geometry and dispersed in Acton Research SPC-500 monochromator after passing through a pump blocking, long wavelength-pass, autofluorescence-free, interference filter (Omega Filters, ALP series). The monochromator was equipped with a CCD camera (Roper Scientific PIXIS-400) allowing for monitoring of the fluorescence spectrum. Fluorescence transients were not deconvolved with the instrument response function since their fluorescent decay lifetimes were much longer than the system response to the excitation pulse. The resulting fluorescent lifetime traces were fit to a single exponential (**eq. 2**)

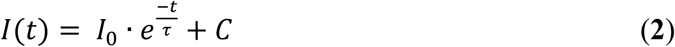

where *I(t)* is the time-dependent intensity, *I_0_* is the initial emission intensity, *t* is time, *τ* is the emission lifetime, and *C* is a constant baseline offset, using MATLAB’s “fittype” function. Standard deviations in *τ* were extracted from the fit using the 68% confidence interval as reported by MATLAB’s “confint” command.

### Preparation of Anaerobic iLOV HQ and OX States

Samples of ∼300 μM purified and reconstituted iLOV or Q_103_X mutants in the flavin OX state were degassed on a Schlenk line before being brought into a COY anaerobic glove box (< 20 ppm O_2_). Simultaneously, solid samples of NaDT and MV^2+^ were brought into the glove box and separately dissolved in anaerobic water to final concentration of 1 M NaDT and 0.1 M MV^2+^. MV^2+^ was quantitated by reduction with excess NaDT and measuring the absorbance at 606 nm of the cation radical (MV•^+^), with an extinction coefficient of 13,700 M^−1^ cm^−1^.^30^ The degassed iLOV OX was then split into two equal volumes. To prepare the HQ iLOV, NaDT and MV^2+^ were added to 100-fold and 10-fold molar excess over iLOV, respectively, to fully reduce the flavin over 15 min. MV^2+^/ MV•^+^ was necessary to kinetically accelerate flavin reduction over NaDT alone, and also served as a visual indicator to ensure NaDT and MV^2+^/ MV•^+^ were subsequently removed during buffer exchange of the iLOV HQ via a Sephadex G25-desalting column.

After reduction, both iLOV OX (unreacted) and HQ samples were buffer exchanged into 20 mM NaCl and 5 mM P_i_ at pH 7.90 via a Sephadex G25-desalting column. The eluted protein samples were then concentrated via Amicon centrifugal filters (10 kDa MWCO) to a volume of about 300 μL before placing them in glass crimp vials and moved to a second VAC Atmospheres glovebox (< 2 ppm O_2_) for further analysis.

In the second glovebox, the OX and HQ protein samples were diluted 20-fold in 200 mM NaCl, 50 mM P_i_ at pH 7.90 and placed in a quartz UV-vis cuvette with a rubber septum. The cuvettes were brought out of the glovebox so that the UV-vis spectra of both the iLOV HQ and OX samples were recorded. The HQ sample was then exposed to air for 2 min with intermittent manual mixing to re-oxidize the sample before re-collecting a spectrum. Concentrations of the initial iLOV HQ and OX samples were then determined by their OX flavin absorbances (448 nm for iLOV OX, 445 nm for Q_103_D OX, or 440 nm for all other mutants). An extinction coefficient of 14,800 M^−1^ cm^−1^ was used for wild-type, while 16,100 M^−1^ cm^−1^ was used for blue-shifted mutants.^31^ Protein samples were used immediately after preparation.

### Anaerobic UV-vis Measurements

UV-vis absorption measurements within a VAC Atmospheres glove box were carried out with the following spectrometer setup, described previously,^32^ with slight modifications. Spectra were collected using a custom fiber optic-connected Ocean Optics QEPro spectrometer and DH-2000-BAL light source. The sample cuvette was mounted in a custom cuvette holder inside the glovebox. The cuvette holder includes fiber optic connectors, a slot for variable sized pinholes at the light-incident face to control light exposure, and a slot for colored glass filters on the light-exiting face (KG-1 or KG-2). The purpose of the pinhole is to narrow the beam of light to reproducibly fit within the confinement of the window of a low-volume 585.3-Q-10 quartz flow cell from Starna Cells. In addition, we have observed that illumination of iLOV OX in anaerobic conditions results in the formation of small amounts of SQ. Thus, the pinhole, in concert with automated shutter control on the light source set only to open when a measurement is being collected, minimizes light exposure and light-induced accumulation of SQ. Control experiments where the frequency of the measurement was doubled showed no effect on SQ accumulation, indicating photochemical SQ formation is indeed minimized by this approach. The use of the flow cell, while not actually being used to flow liquids during measurements, allows for very small sample volumes (70-80 μL) to be used. The purpose of the glass filter is to improve the balance of the light source output across the region of the spectrum where OX and SQ absorb (450‒650 nm), improving the signal-to-noise ratio across this region. A buffer blank was used to establish zero absorbance and a linear baseline correction was used by subtracting a least-squares linear regression fit calculated between 680 and 900 nm—where no species exhibits absorption—and extrapolated across the entire wavelength range. Linear baseline shifts were observed with even minor mechanical adjustments to the fiber cables, but otherwise does not affect the spectrum.

### Measurement of NSQ Molar Absorptivity

To quantify NSQ formation by comproportionation (*vida infra*) we measured the extinction coefficient (ε) of the NSQ at its maximal absorbance (λ_max_) at the red-shifted absorption feature at 616 nm (ε_616_). The iLOV Q_103_D mutant is light-sensitive and undergoes quantitative photo-reduction to the NSQ state when exposed to 450 nm light, making it an ideal candidate for estimation of the ε_616_, which was assumed to be identical across the iLOV variants in this study.^33^ Samples of OX Q_103_D were brought into a VAC glovebox after degassing, as described above. A quartz cuvette with an open top was prepared containing 400 μL of roughly 25 μM protein in a buffer composed of 200 mM NaCl and 100 mM MES at pH 6.50. As the sample sat in a cuvette holder, it was illuminated directly from above with a 2 W 450 nm LED, and photoreduction was monitored spectrophotometrically. The increase in sample absorbance at 616 nm indicated the generation of NSQ, which reached equilibrium after 10 minutes (**Supplemental Information Figure S2**). At this point the LED was turned off to remove interference from the excitation light, and the final spectrum of the semiquinone species was obtained. The absorptivity of the NSQ was estimated from the ratio of the absorbance at 445 nm (which we assumed had an absorptivity of 16,100 M^−1^ cm^−1^ from the similarly blue-shifted Q_103_K mutant with λ_max_ of 440 nm)^31^ and the absorbance at 616 nm with ε_616_ of 8,200 ± 100 M^‒1^ cm^‒1^.

### Electron Paramagnetic Resonance (EPR)

As the value of ΔE was being measured optically for each mutant, an EPR sample was produced simultaneously with equimolar quantities of OX and HQ between 200 and 400 μM total protein and a final volume of 300 μL with 10% glycerol and was kept in the dark. When the low concentration samples monitored by UV-vis spectroscopy reached equilibrium, the EPR sample was placed into a high-precision quartz EPR tube and flash frozen in liquid N_2_-cooled isopentane (< −130 °C) cold well within the glovebox. The sample was then removed from the glovebox and placed into a storage dewar in liquid N_2_ before analysis by X-band EPR spectroscopy on a Bruker EMXplus EPR spectrometer. Because the Q_103_D iLOV variant has the least crossed potentials the radical concentrations were large enough to be quantified and measured in triplicate. X-band EPR and radical quantitation was performed as described previously.^32^ The spectra were simulated using EasySpin 6.0 in MatLab, and spin quantitation was performed by double integration of the EPR signal, divided by the Q-value, and comparison to three 4-hydroxy-TEMPO standards. The concentration of the 4-hydroxy-TEMPO standards were determined by the absorbance of a stock solution using the reported extinction coefficient at 428 nm of 12.4 M^−1^ cm^−1^.^34^

### Purifying and Characterizing Isomers of Safranine

Commercially available safranine O contains several isomers with distinct optical and redox properties.^35^ To ensure reliable redox titrations with a dye indicator of a known reduction potential and extinction coefficient, we purified the isomers of commercially available safranine O using a Shimadzu Nexera preparative high-performance liquid chromatography (HPLC) system with an inline single quadrupole mass spectrometer (MS) detector and UV-vis detector. The isomers were resolved on a 19 mm x 150 mm XBridge BEH C18 OBD Column with 130 Å pore size, 5 μm particle size, using a stepwise elution gradient, with acetonitrile concentrations increasing from 12.5% to 13.5% over 25 minutes, and then 13.5% to 20% over 2 min. Three different species were separated, labelled “1”, “2”, and “3”. Isomer 1 was not sufficiently resolved from isomer 2 to be used, but isomers 2 and 3 fractions were sufficiently pure for analysis and use. Both isomers were analyzed by ^1^H NMR in *d*_6_-DMSO on a 500 MHz Bruker Avance NEO spectrometer. Amber NMR tubes were used to protect the sample from light. Structural assignments of isomer 2 and 3 are reported in the **Supplemental Information Figure S4** and **Table S3**. The reduction potential of isomer 3 (1 mM) in phosphate-buffered saline (PBS; 137 mM NaCl, 2.7 mM KCl, 10 mM Na_2_PO_4_, 18 mM K_2_PO_4_) was measured as a function of pH by cyclic voltammetry with a pyrolytic graphite (edge plane) working electrode, a platinum wire counter electrode, and a Ag/AgCl (3 M NaCl) reference electrode (**Supplemental Information Figure S5**). Potentials were scanned from −0.2 V to −1 V with a scan speed of 100 mV/s. The Ag/AgCl reference electrode potential was calibrated against a saturated calomel electrode (SCE) using a multimeter and the resulting potentials were adjusted to be relative to the standard hydrogen electrode (SHE). Based on the close match in E° of the safranine isomer 3 and iLOV, this isomer was used for all redox titration experiments and is simply referred to as “safranine,” henceforth.

### Measurement of iLOV E°_OX/HQ_

To measure the equilibrium two-electron E°_OX/HQ_ potential, we developed a method to perform redox titrations using a 96-well plate format and a BMG VANTAstar microplate reader housed inside a Coy anaerobic chamber (< 20 ppm O_2_). To prepare the plate, a master mix containing 3× Buffer A (600 mM NaCl, 150 mM P_i_ at pH 7.9) or 3× Buffer B (600 mM NaCl, 300 mM Bicine at pH 8.9), 180 μM iLOV_OX_, 90 μM of MV^2+^, and 240 μM safranine were aliquoted into several wells of a 96-well plate at a volume of 50 μL. Different ratios of water and a 700 μM NaDT stock were combined into each well to bring the final volume to 150 μL. At least one well contained no reductant, and one well contained excess (3.3-fold molar equivalents) reductant to serve as reference spectra for 100% OX and 100% HQ respectively. In wells in which excess NaDT was added, an authentic spectrum of MV^+^• mediator was subtracted from the resultant spectra using the absorbance at 606 nm. In these wells with low solution potential, it can be assumed that the concentration of SQ is too low to interfere with the absorbance from MV^+^• at this wavelength. A typical plate measurement contained 14 different reducing conditions, 3 different replicates, and 2 different pH conditions using a total of 84 wells. Remaining wells were used for blanks and standard spectra of methyl viologen in 1× Buffer A to be subtracted during the analysis. The 96-well plate was then sealed and placed into the plate reader, and spectra from 300 to 900 nm with a resolution of 5 nm were collected from each well, averaging a total of 22 flashes each, every 30 min for 5 to 12 h, though samples often reached equilibrium after about 5 h.

The redox titration UV-vis data was analyzed similarly to previous treatments,^36,37^ with some notable differences. The proton-coupled half-cell reactions for the unknown analyte (iLOV) and the standard indicator dye (safranine) are represented by reactions **1** and **2**,

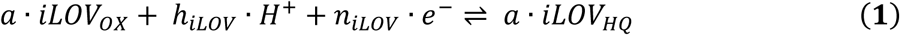

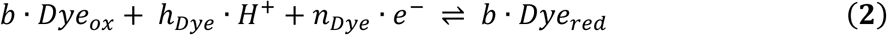

where a, b, h, and n are the stoichiometric coefficients for iLOV, safranine, H^+^, and *e*^‒^, respectively. These two reactions can be scaled by dividing reaction **1** by *n_iLOV_* and multiplying by *n_dye_/b* and dividing reaction **2** by *b*, such that the same number of electrons transferred in both reactions are equal via reactions **3** and **4**.

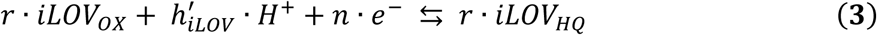

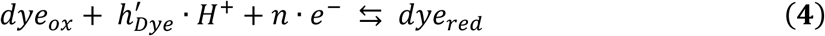

Here, the new stoichiometric coefficients, *r*, *h′_iLOV_*, *h′_dye_*, and *n* are related to the original stoichiometric coefficients by r = (*a/n_iLOV_*)/(*b/n_dye_*), *h′_iLOV_* = (*h_iLOV_*/*n_iLOV_*)/(*b/n_dye_*), *h′_dye_* = (*h_dye_*/*b*), and *n* = *n_dye_/b*. If the two reactions are in equilibrium, within the same reaction vessel at the same pH, and assuming an activity coefficient of unity, then the standard cell potential for each half-reaction (*E°*) is related to the solution potential (*E*_sol_) by **eq. 3** and **4**.

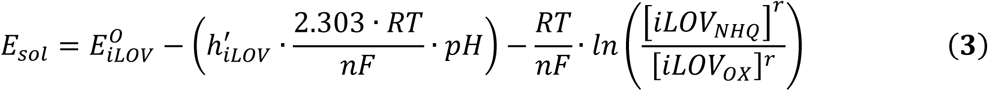

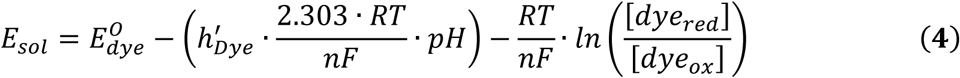

At a given point in the titration, *E_sol_* dictates the ratio oxidized and reduced states of the standard dye indicator (safranine) and the unknown analyte (iLOV) based on their respective *E°*. Setting these equations equal to one another at *T* = 298 K, and *n_iLOV_/a* = *n_dye_/b* = 2, such that *r* = 1, *n* = 2, and *h′_iLOV_* = *h_iLOV_/a*, and then rearranging into a linear format, we obtain **eq. 5**.

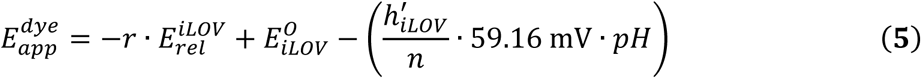

where *E^dye^ _app_* and *E^iLOV^ _rel_* are defined by **eq. 6** and **7**.

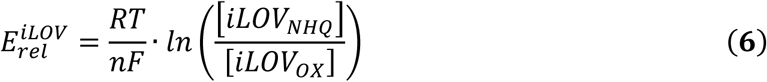

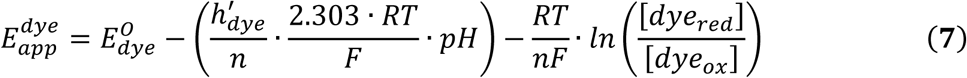

We determined the pH dependent reduction potential of safranine *E^dye^_app_* directly from the Pourbaix diagram in **Supplemental Figure S4**. Thus, linear plots of *E^dye^_app_* vs. *E^iLOV^ _rel_*, generated by spectrophotometric measurement of *ln([dye_red_]/[dye_ox_])* and *ln([iLOV_HQ_]/[iLOV_OX_])*, one obtains a slope of *r* (1 for stoichiometric reduction of dye and iLOV by two electrons), and a y-intercept equal to the *E°_OX/NHQ_* of iLOV at a given pH.

Using the absorption of iLOV FMN_OX_ on the blue edge of the S_0_→S_1_ absorption band (420 nm) to minimize interference from safranine, and 520 nm for oxidized safranine, the concentration of the oxidized species was determined and converted to fraction oxidized and fraction reduced to calculate *ln([x_red_]_frac_/[x_ox_]_frac_)*, where *[x_ox_]_frac_ = [x_ox_]_x_/[x_ox_]_i_*, and *[x_red_]_frac_ = 1 ‒ [x_ox_]_frac_*. These relationships hold for situations where both the iLOV and safranine equilibria are composed of only the fully reduced or oxidized states, simplifying the analysis. However, in the Q_103_K and Q_103_D iLOV, appreciable SQ accumulates during these titrations, and must also be accounted for, but the E°_OX/HQ_ midpoint potential for OX/HQ is still defined as the solution potential at which the concentration of OX and HQ are equal. To quantitate SQ contributions to the iLOV population during redox titrations, and therefore accurately estimate HQ, the concentration of NSQ was estimated using the absorbance at 616 nm and the aforementioned extinction coefficient of 8,200 M^−1^ cm^−1^ for each well. Next, an authentic NSQ spectrum collected as described above, was scaled to the absorbance at 616 nm and subtracted from each spectrum to obtain the concentration of OX at 420 nm without convolution with NSQ. From this, [x_red_]_frac_ = 1 ‒ [x_ox_]_frac_ ‒ [x_nsq_]_frac_ was used to estimate HQ. Data points in the final plot outside the range of 0.05 < [x]_frac_ < 0.95 were ignored in the linear regression. The error at each point was determined from the standard deviation of triplicate measurements. Replicates with obvious aberrations (such as resulting from a bubble within a well) were ignored as outliers.

### Comproportionation of iLOV HQ and OX

When equimolar concentrations of OX and HQ are mixed (that is, initially, [OX]_i_ = [HQ]_i_), they will comproportionate to an equilibrium concentration of [SQ]_eq_, [OX]_eq_, and [HQ]_eq_ via reaction **5**,

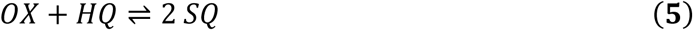

with an associated equilibrium constant (*K*_eq_, **eq. 8**),

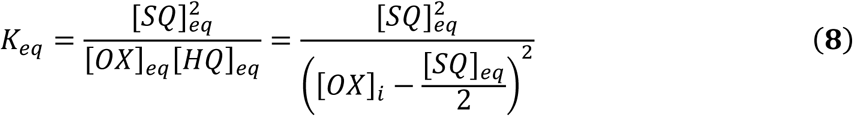

where *[X]_eq_* is the equilibrium concentration and *[X]_i_* is the initial concentration of species *X*. **Eq. 8** is equivalent to the product of the *K*_eq_’s for its respective one-electron half reactions **6** and **7**,

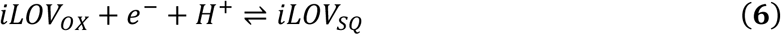

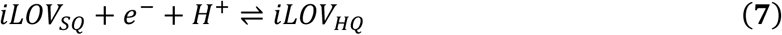

meaning that the Nernstian potential derived from *K*_eq_ corresponds to difference between the two one-electron couples, E°_OX/SQ_ and E°_SQ/HQ_, by **eq. 9**,

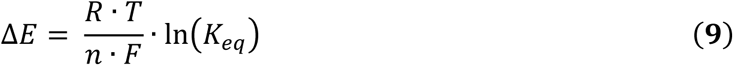

where *ΔE* is the relative electrochemical potential separation of E°_OX/SQ_ and E°_SQ/HQ_ couples, *R* is the gas constant, *T* is the temperature in Kelvin, *n* is the number of electrons (1 for the comproportionation reaction), and *F* is Faraday’s constant. In our convention, *ΔE* is equivalent to the extent of potential “crossing”, where negative values are crossed, and positive values are “normally-ordered” potentials. If the absolute midpoint potential for the two-electron process (E°_OX/NHQ_) is known, it becomes straightforward to calculate the values of the two one-electron reduction potentials from ΔE by **eq. 10** and **11**.

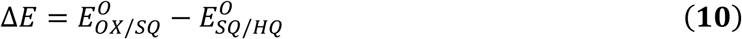

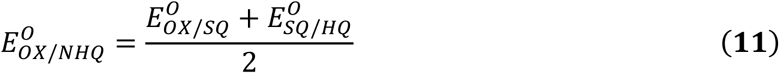

To measure ΔE, pure samples of iLOV or Q_103_X mutants in the OX and HQ state were prepared and quantified as described earlier inside a VAC Atmospheres glove box (< 0.2 ppm). These two species were combined to a final concentration of 50 μM of OX and HQ in 200 mM NaCl and 50 mM P_i_ at pH 7.90, mixed, then quickly loaded into the flow cell and placed into the cuvette holder for a kinetics measurement. About 10-15 s passed between mixing the sample and beginning the kinetics measurement. The resulting time-dependent increase in NSQ concentration was monitored by the absorbance at 616 nm until it reached equilibrium (5-24 h), fit to a single exponential, and the exponential function pre-factor was used to quantify NSQ based on the ε_616_ of 8,200 M^−1^ cm^−1^. The value of *K*_eq_ was then calculated using **eq. 8** with [iLOV_OX_]_i_ = 50 μM and ΔE was extracted from **eq. 10**. Values of E°_OX/NSQ_ and E°_NSQ/NHQ_ were obtained by respectively adding and subtracting ΔE/2 from the previously measured value of E_OX/HQ_. Each mutant was measured in triplicate at pH 7.90.

### Molecular Dynamics, Hybrid Quantum Mechanical/Molecular Mechanical (QM/MM), and Alchemical Free Energy Calculations

We conducted molecular dynamics (MD) simulations of iLOV and Q_103_X mutants in the OX and NSQ states. The force field parameters for the ribityl phosphate tail were obtained from the Manchester AMBER parameter database,^38^ and the parameters for the isoalloxazine ring in the OX and NSQ state were parameterized with Antechamber using the restrained electrostatic potential (RESP).^39^ Force constants for bonds, angles, and torsions were taken from the GAFF parameter set.^40^ To set up the model, we centered the protein and cofactor (flavin) in a cubic box at 2 nm from the edges and solvated with TIP3P waters.^41^ Na^+^ ions were added to neutralize the system. Additional Na^+^ and Cl^-^ ions were added to mimic the experimental salt concentration (0.2 M). MD simulations were performed using GROMACS 2024.4 software package along with amber99sb force field.^42^ Bonds involving hydrogen atoms were constrained using LINCS algorithm,^43^ and 2 fs timestep was used. The van der Waals interactions were cutoff at 0.8 nm. All the trajectories were carried out in the NPT ensemble with the C-rescale barostat with τₚ = 5.0 ps and τₜ = 2.0 ps with the V-rescale thermostat. Energy minimization was performed in three stages, 1) solvent and ions only, 2) solvent, ions, and protein side chains without hydrogen atoms, 3) all atoms except the FMN cofactor. The system was then equilibrated for 80 ns in two phases: a 0.3 ns heating step from 0 K to 298 K at 1 atm, followed by 79.7 ns equilibration. These equilibration trajectories were discarded. Finally, NPT production simulations of 500 ns were performed for both OX and NSQ states for subsequent analysis. A similar procedure was followed to simulate 500 ns trajectories for each of iLOV’s Q_103_ mutations—arginine (R^(+)^), aspartic acid (D^(0)^), glycine (G), alanine (A), and lysine (K^(+)^). The mutations were introduced to the model using SCWRL4.0.^44^

Using the 500 ns MD simulations described above, we further conducted statistical clustering analysis. The trajectories were aligned on the Cα backbone atoms and clustered for the whole system with the GROMOS algorithm with a 0.2 nm root mean square displacement (RMSD) cut-off.^45^ This choice of cut-off distance was reached by testing the effect of the cut-off value on the population of the largest cluster for parental iLOV. The results of this testing are shown in **Supplemental Information Figure S3**. The optimal choice was a cutoff that produces only a few clusters (around 5-10), and where the largest cluster captures a significant percentage of the population (66% in parental iLOV).

Root mean square fluctuation (RMSF) values were computed for the Cα atoms of the wild-type protein in both the oxidized (OX) and neutral semiquinone (NSQ) states as well as for selected mutants using MDAnalysis.^46^ The results were visualized as a residue-by-state heatmap.

Water dynamics around the N5 atom of the flavin cofactor were analyzed using a custom MDAnalysis-based script. Trajectories and topologies (xtc and tpr) were loaded, and the protein backbone was aligned to remove global translation and rotation. For each frame, the closest water oxygen to N5 was identified, and the following metrics were recorded: N5–water distance, hydrogen-bond formation (distance ≤ 3.5 Å, angle ≥ 150°), and whether the water occupied a defined pocket (within 4 Å of N5 or O4′ were recorded; this parameter indicates presence of water nearby but one that is not necessarily in the right orientation to form a hydrogen bond).

The *gmx hbond* tool was used to determine the total number of H-bonds formed by each of the N1, O2′, N3, O4′, and N5 heteroatoms of flavin with the surrounding water and protein environments in the parental iLOV and its mutants in each of the OX and NSQ states.

To simulate the effect of a mutation on the reduction potential of iLOV, a thermodynamic cycle with alchemical transformation of Q_103_ to various mutants was used (**Figure 2**).^47,48^ The free energy associated with alchemical mutagenesis of iLOV’s Q_103_ residue to the mutant for each of the OX and NSQ states provided two legs of the thermodynamic cycle, while the third leg, corresponding to the free energy of reduction of flavin in the wild-type system, is obtained from the experimental data (*vida infra,* ‒458 mV or 44.19 kJ/mol). Using those three quantities, the fourth leg of the thermodynamic cycle was computed to provide the free energy of reducing flavin in the mutated iLOV. This free energy is then converted to a reduction potential using **eq. 12.**

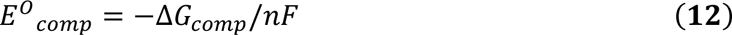

**Figure 2.**
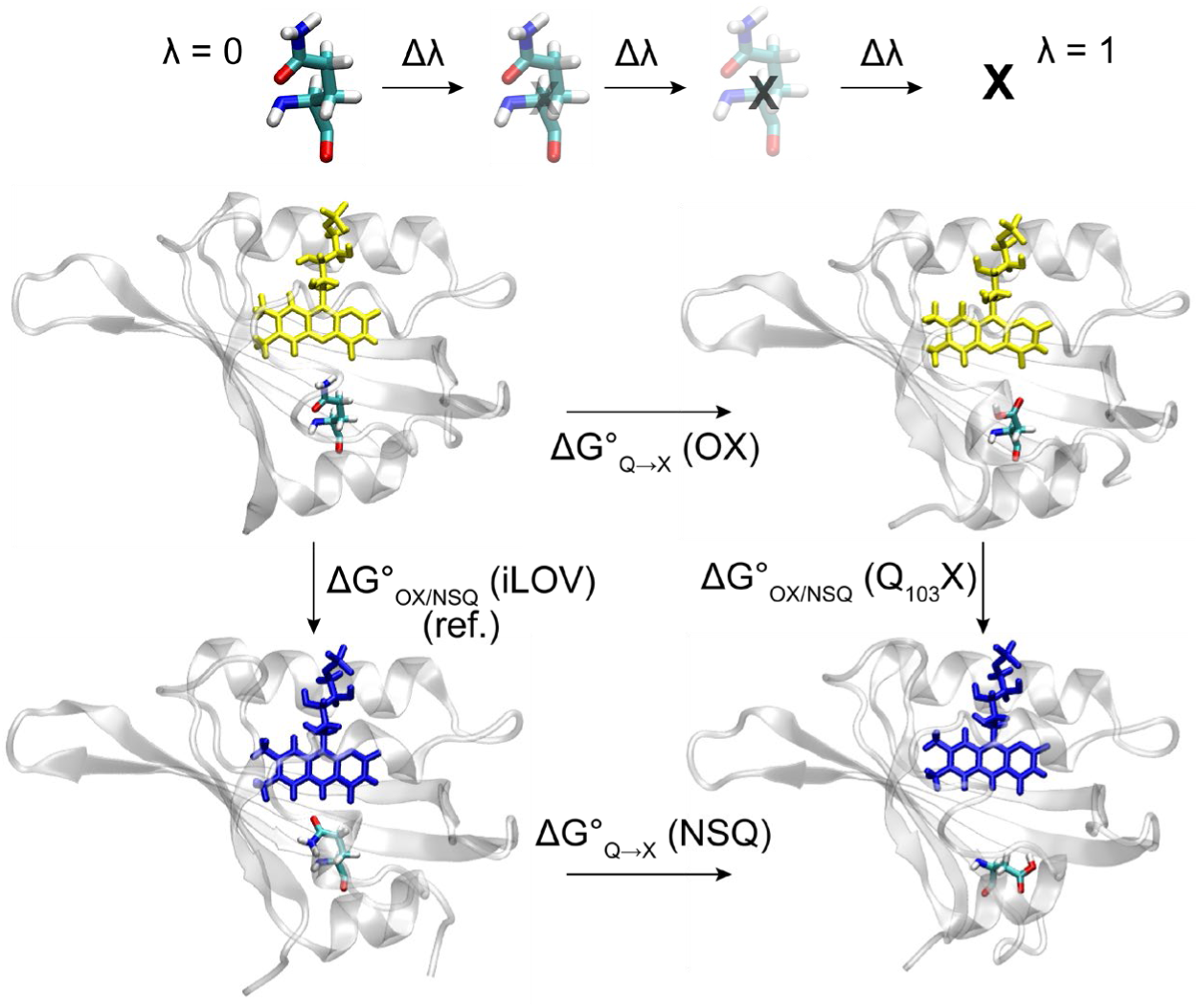
Thermodynamic cycle used to calculate ΔG°_OX/NSQ_, and hence E°, for Q_103_X mutants via alchemical transformations.

The Bennett Acceptance Ratio (BAR) approach was used to compute the free energy of the alchemical transformation steps.^49–51^ For this purpose, we adopted a protocol that builds on the recently automated Average Protein Electrostatic Configuration QM/MM approach for flavoproteins (APEC-F 2.0).^52^ A similar idea, albeit with several modifications in the details of the protocol, was used to simulate relative free energies and isomerization barriers for retinal binding proteins and molecular switches in solution.^53^ Briefly, with APEC-F 2.0, QM/MM models of iLOV in its oxidized (OX) and neutral semiquinone (NSQ) states were constructed starting from the X-ray crystallographic structure (PDB ID 4EES).^24^ We focus on the OX/NSQ simulations because these are the ones displaying the strongest variations with mutations experimentally. Using APEC-F 2.0, the charges and geometry of the flavin optimized quantum mechanically were computed at the B3LYP/ANO-L-VDZP level of theory in a superposition of 100 solvated protein configurations extracted from molecular dynamics (MD). As done for the 500 ns MD simulations, the system was neutralized using a 0.2 M salt concentration. Details of the QM/MM model construction and methodology were reported recently.^52^ Software employed in the construction of the APEC-F 2.0 models include OpenMolcas, Tinker, GROMACS, and Dowser.^50,54–56^

The APEC-F 2.0 QM/MM structure provides a starting point for further free energy simulations computed with BAR as implemented in the GROMACS 2024.4 *gmx bar* tool.^50,51^ Since the redox site on flavin is localized on the rigid isoalloxazine moiety, the cofactor was kept frozen in its QM/MM-optimized geometry throughout the entire BAR protocol, while relaxing the protein, TIP3P water molecules, and solution ions during the MD. The protocol was run in the NVT ensemble using the equilibrated volume that was originally determined at the level of APEC using NPT dynamics. Hydrogen bonds were treated using LINCS^43^ and MD simulations used a timestep of 2 fs. Long-range electrostatic interactions were treated with Particle Mesh Ewald (PME),^57^ with a real space cut-off of 0.8 nm, Fourier grid spacing of 0.12 nm, PME order of 4, and relative strength of the cutoff at 10^‒5^. The van der Waals interactions were cut off at 0.8 nm. The alchemical mutations were introduced using the pmx library adopting the amber99sbmut force field,^58^ which includes dummy atoms in a dual topology fashion. The alchemical transformation was carried out in a series of 40 discrete λ windows connecting the two physical end points: the parental iLOV protein (λ = 0) and mutated amino acid (λ = 1), as shown in Figure 2. We adopted a serial equilibration approach where the last frame of equilibration at a specific λ step is the starting frame for the next one. Specifically, for each λ window, we equilibrated the system (protein + cofactor) for 3 ns in NVT followed by 12 ns production runs, giving a total simulation time of 600 ns for each replica. For each mutant and each of the OX and NSQ states, we ran multiple (≥3) replicas. A few replicas were also carried out using longer initial equilibration (50 or 100 ns) at the λ=0 step to serve as an additional reference and reduce uncertainty in the results when needed.

Redox potential shifts were computed in this way for mutating Q_103_ in iLOV. For mutations involving the introduction of a charge (Q_103_D^(‒)^, Q_103_K^(+)^, and Q_103_R^(+)^), we adopt the co-alchemical ion approach where an ion is co-perturbed into existence along with the mutant at each λ to keep the net charge of the system constant.^59^ Specifically, for Q_103_D^(‒)^, a growing negative charge arising in the simulation as the glutamine is annihilated and aspartate forms was neutralized with the gradual morphing of a water molecule into a sodium ion. Similarly, for Q_103_K^(+)^ and Q_103_R^(+)^, the introduced positive charge was neutralized by morphing a water into a chloride ion.

To monitor water leakage into the flavin binding site along the BAR perturbation steps, the *gmx rdf* tool was used to determine the radial distribution function (RDF) of water relative to flavin’s N5 along each λ trajectory,^60^ as well as the cumulative number (CN) obtained by integrating the RDF. The CN value represents the average number of particles within a given distance.

## RESULTS

During flavin reduction, N5 becomes protonated during the OX→NSQ transition while N1 becomes protonated during the NSQ→NHQ transition, depending on pH and the flavin environment, which modulates the N5 and N1 p*K*_a_s.^7,61^ We hypothesized that hydrogen bonding interactions at the N5 and N1 sites could confer independent mechanisms of control of the two one-electron couples in flavoproteins via thermodynamic control of proton-coupled electron transfer (PCET). In this work, we focused on N5 using the small flavoprotein, iLOV, as a model system.

The X-ray crystal structure of iLOV reveals a glutamine residue (Q_103_, **Figure 3A**),^24^ positioned near both N5 (3.4 Å) and O4′ (3.1 Å) of the bound FMN cofactor. To investigate how residues near N5 and O4′ of a flavin cofactor affect the flavin reduction potentials, we generated Q_103_X mutations (X = alanine, A; glycine, G; arginine, R; lysine, K; and aspartic acid D) by site-directed mutagenesis. All purified proteins quantitatively bound FMN at a 1:1 ratio during the reconstitution (**Supplemental Information Tables S1** and **S2** and **Figure S1**).

**Figure 3.**
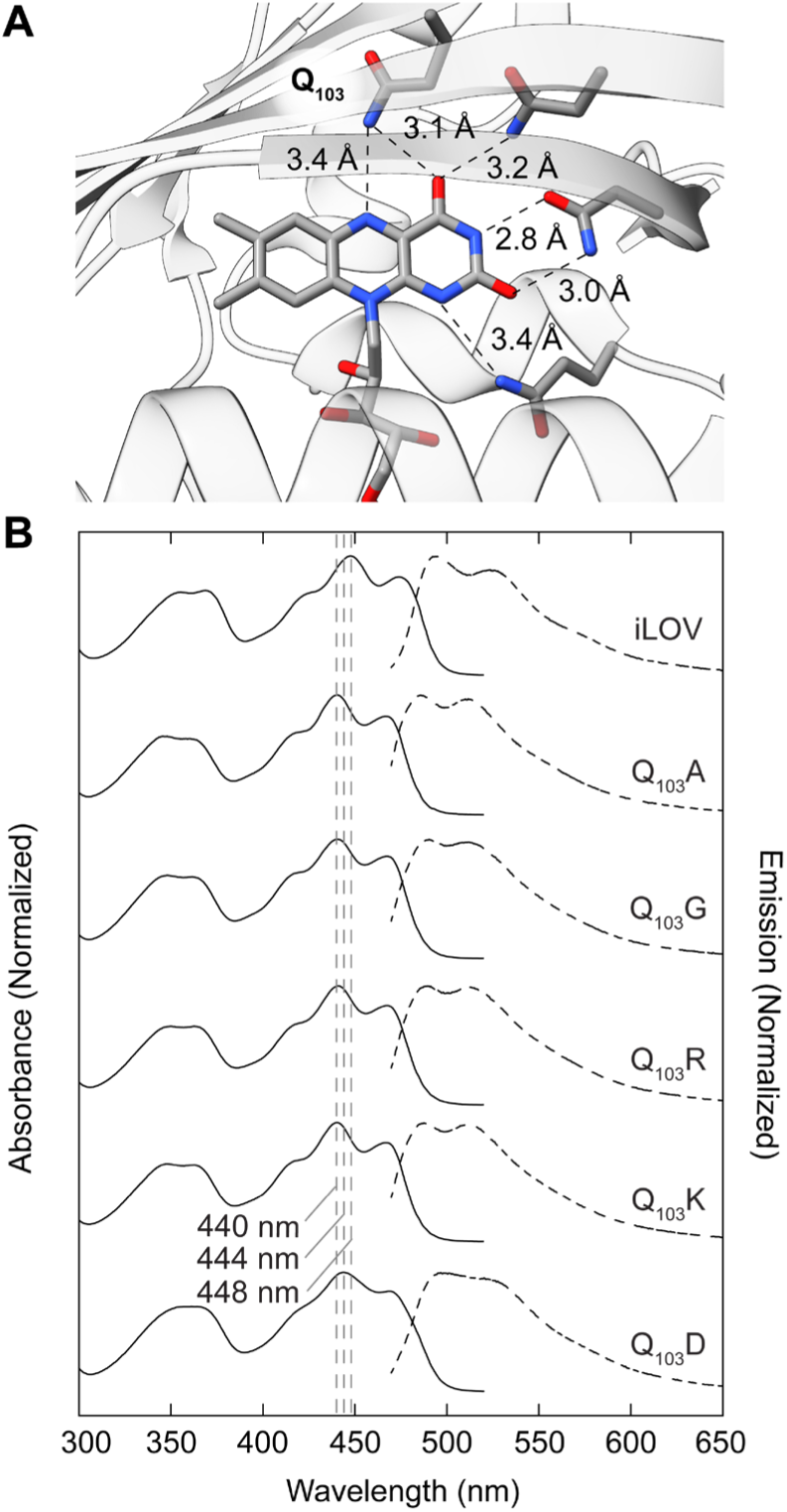
Hydrogen bonding interactions to N5 in iLOV and effect of Q_103_X mutations on optical properties. **A** Interactions of iLOV FMN cofactor (PDB ID: 4EES) with the protein environment. **B** Normalized UV-vis absorption and emission spectra (λ_ex_ = 450 nm) of iLOV and Q_103_X mutants.

The absorption spectrum of the OX state of the cofactor was modestly perturbed by the Q_103_X substitutions; the parent iLOV exhibited a λ_max_ of the S_0_ → S_1_ transition at 448 nm, whereas the Q_103_D λ_max_ was blue-shifted to 444 nm, and Q_103_A, Q_103_G, Q_103_K, and Q_103_R were further blue-shifted to 440 nm (**Figure 3B**). Emission and excitation spectra, emission λ_max_, quantum yields (Φ), and associated emission lifetimes (τ) were also modestly changed by the Q_103_X mutations (**Supplementary Information Figure S6**, **Figure S7**, and **Table S4**). For iLOV, the emission λ_max_ is at 492 nm with a Φ of 0.37 ± 0.01. All of the Q_103_X mutants had blue-shifted λ_max_ between 2 nm to 6 nm, generally consistent with their blue-shifted absorption λ_max_, with the exception of Q_103_D, which was red-shifted to 498 nm. The Q_103_X mutants exhibited similar Φ, ranging from 0.37 ± 0.01 in Q_103_A to 0.33 ± 0.01 in Q_103_D. The fluorescence emission lifetime for iLOV was 4.37 ± 0.01 ns, while mutations to Q_103_ resulted in a span from 4.01 ± 0.01 for Q_103_A to 4.11 ± 0.01 for Q_103_R. Interestingly, τ did not correlate with measured Φ, a phenomenon that has been observed when placing certain fluorophores in solvents of varying polarity.^62^ This is thought to result from simultaneous changes to both the radiative and non-radiative relaxation rates, which we expect are sensitive to the N5 environment in iLOV and the associated Q_103_X mutants.^63^

We next examined how Q_103_ mutation affected the redox behavior of the flavin cofactor, given its proximity to N5, its hydrogen bonding character, and the change in hydrogen bonding character of N5 during redox cycling. Our initial attempts using protein film voltammetry (PFV) and the xanthine/xanthine oxidase (XXO) method were inconclusive. Although iLOV was capable of adsorbing to a pyrolytic graphite edge plane electrode, the electrochemical responses were variable, with unexpected peak current separation and amplitudes, and showed no differences between the parent iLOV and Q_103_X mutants (**Supplementary Information Figure S8**). The XXO assay was also unsuitable due to slow electron transfer kinetics (**Supplementary Information Figure S9**). We, therefore, turned to redox titrations, using safranine as a dye redox standard reporter and NaDT as a reductant at pH 7.9 and 8.9 (**Figure 4**). Upon addition of the reductant, we observed a loss of the characteristic visible absorbance bands of the OX FMN (448 nm) and the oxidized safranine dye (520 nm, **Figure 4A**), that disproportionately reduces safranine before equilibrium is reached between safranine and iLOV based on their relative reduction potentials (**Figure 4A**, inset). After two equivalents of NaDT, iLOV and safranine were completely reduced, and no clear accumulation of an NSQ or ASQ intermediate was observed at the titration midpoint at either pH. This is consistent with significantly crossed potentials and an unstable SQ, as reported previously.^64^

**Figure 4.**
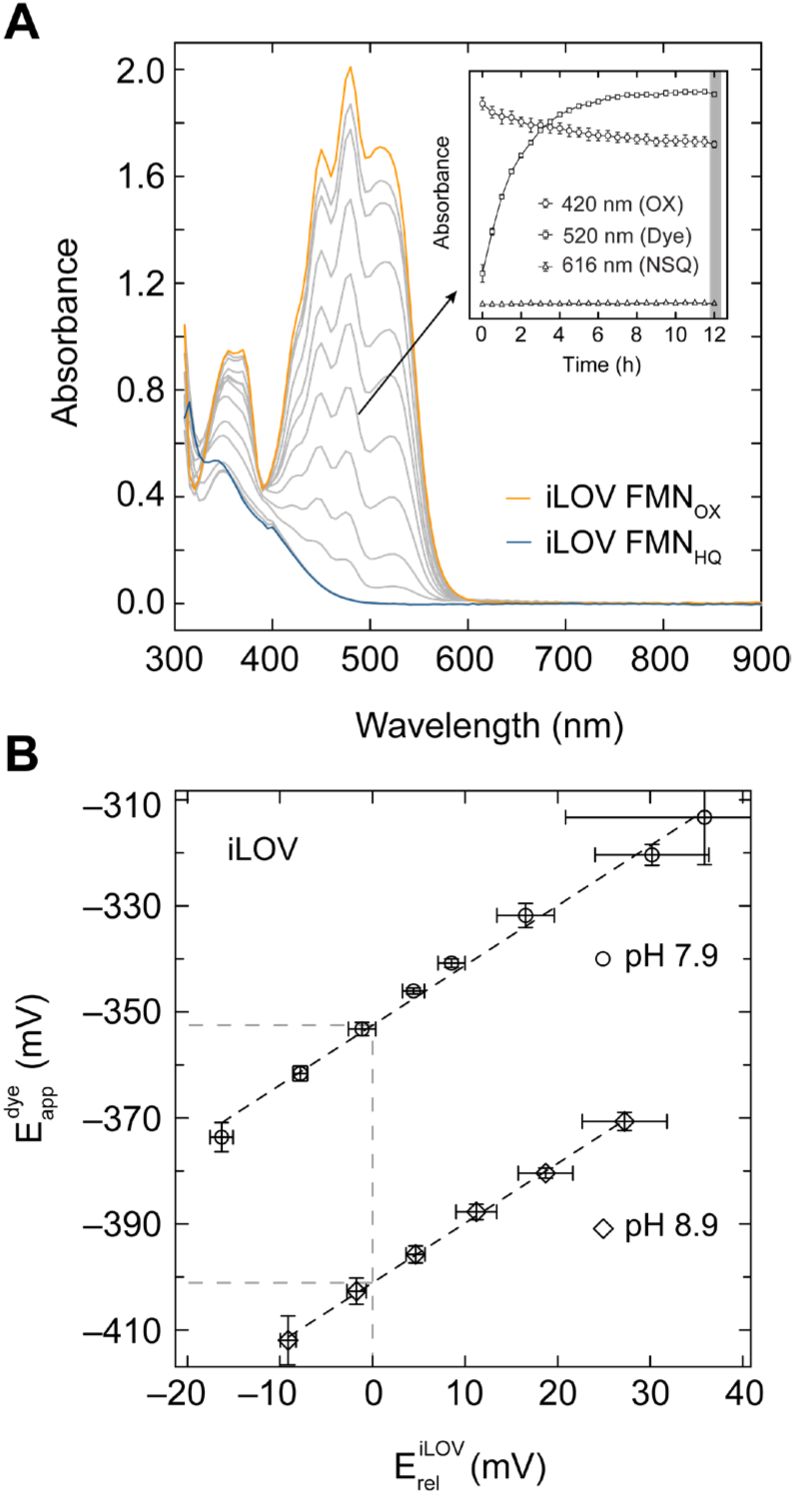
Spectrophotometric redox titrations of iLOV. **A** Equilibrium iLOV absorbance spectra after adding NaDT reductant from 0 eq (orange) to 3.3 eq. (blue). Inset shows the approach to equilibrium at one point in the titration (arrow) via monitoring the absorption at 420 nm (FMN_OX_), 520 nm (safranine), and 616 nm (FMN_NSQ_). Safranine is initially reduced faster than iLOV, then reoxidized by reducing iLOV. **B** E_app_ (determined from safranine) vs. E_rel_ (determined from iLOV) plots for pH 7.9 (blue circles) and pH 8.9 (orange circles) and associated linear fits. The gray dashed line shows the E_app_ at which the iLOV FMN has an OX/HQ ratio of 1 (*i.e.* E°_OX/NHQ_).

Assuming an equilibrium dominated by OX and HQ forms of the flavin, the fractional reduction of both iLOV Q_103_X and the safranine reporter dye standard were determined and their natural logarithms were plotted against one another, yielding a linear relationship, where the slope indicates the ratio of the stoichiometric coefficients for the number of electrons transferred between the protein and the indicator dye, the y-intercept reflects the E°_OX/HQ_ determined relative to the standard, and the pH dependence reflects the number of protons involved in the redox process (**Figure 4B** and **Supplemental Information Figure S10**). As the apparent solution potential from the dye indicator (E^dye^_app_) decreases, so too does the relative potential of iLOV, (E^iLOV^_rel_), and the point at which the concentration of the OX and HQ states are equal we determine the E°_OX/HQ_ (**Figure 4B** dashed lines). The resulting extracted values for E°_OX/HQ_ for iLOV and the Q_103_X mutants, as well as the reference values for free FMN are summarized in **Table 1**.

**Table 1.**
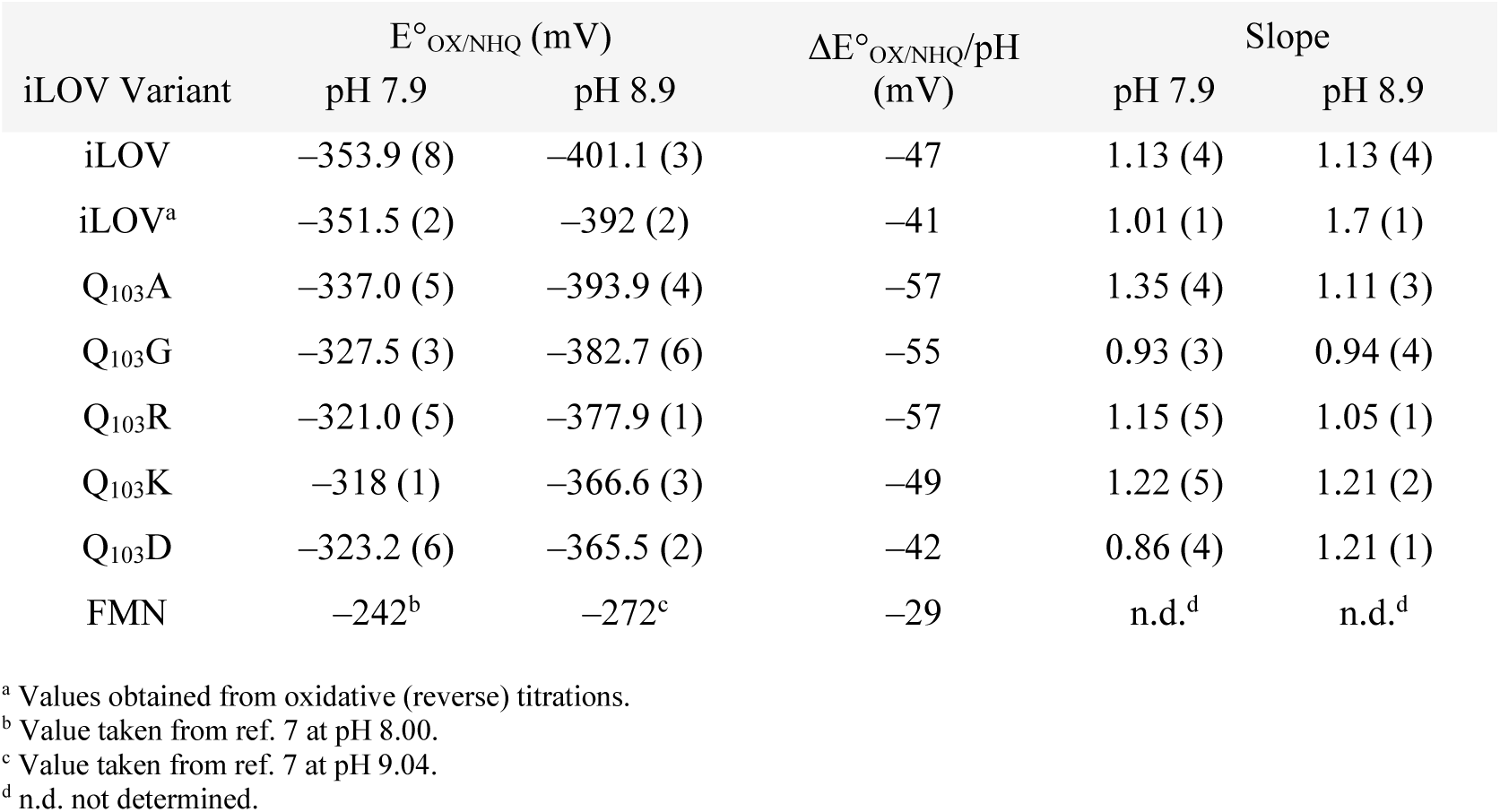
Reduction potentials, pH dependence, and electron equivalents with the mediator (Slope) of iLOV and Q103X mutants determined from reductive and oxidative (iLOV) redox titrations. Uncertainties in the last significant digit shown in parentheses.

From the redox titrations, the parent iLOV had the lowest E°_OX/HQ_ of −353.9 ± 0.8 mV at pH 7.9 and a pH dependence between pH 7.9 and 8.9 of ‒47 mV/pH, lower than the theoretical value for a 2 *e*^‒^/2 H^+^ couple of ‒59 mV/pH, but substantially higher than the expected ‒29 mV/pH for a 2 *e*^‒^/1 H^+^ couple. These data are most consistent with iLOV reduction proceeding via concerted OX→NSQ and NSQ→NHQ transitions. The reaction was fully reversible; an oxidative titration of reduced iLOV and safranine resulted in an E°_OX/NHQ_ potential only 2.5 mV different, with slightly larger differences at pH 8.9 (9 mV), but significantly more scatter in the data (**Supplemental Information Figure S11**). Extrapolation to pH 7.0 yields a E°′_OX/NHQ_ of ‒312 mV, approximately 30 mV lower than that measured for the *Chlamydomonas reinhardtii* LOV1 C_57_G protein,^64^ and 41 mV lower than the homologous *Avena sativa* LOV2.^65^

The mutants Q_103_A, Q_103_G, and Q_103_R also showed no significant SQ accumulation during the redox titrations. The E°_OX/NHQ_ followed the trend iLOV < Q_103_A < Q_103_G < Q_103_R and an average ΔE°_OX/NHQ_/pH of ‒56 mV/pH, with the highest E°_OX/NHQ_ shifted +33 mV relative to the parent iLOV. During the titrations of both Q_103_K and Q_103_D, we did observe some amount of NSQ, based on its characteristic absorbance at 616 nm, with greater NSQ accumulation for the Q_103_D mutant relative to Q_103_K. In both mutants, the NSQ accumulation was not quantitative, suggesting E°_OX/NSQ_ and E°_NSQ/NHQ_ were only moderately separated. Because the SQ contribution complicated two-state analysis of E°_OX/NHQ_, we quantified NSQ directly and accounted for it in the calculation of E°_OX/NHQ_. Taking advantage of the unique photosensitivity of the Q_103_D,^33^ we quantitatively photo-reduced the OX FMN to the NSQ state anaerobically via 450 nm illumination to determine the molar absorptivity at 616 nm (ε_616_) (**Supplemental Information Figure S2**). This reaction requires no external reductant; presumably the electron is donated by a protein tyrosine or tryptophan. For further validation and to capture any contributions of an ASQ, we also quantitated the total SQ population by EPR, against a 4-hydroxy-TEMPO standard (**Supplementary Information Figure S12**). The ε_616_ determined from these EPR measurements is 12,400 ± 400 M^−1^ cm^−1^, compared to our estimated value of 8,200 ± 100 M^−1^ cm^−1^ by UV-vis. Due to the challenges of EPR quantitation by double integration, we use the UV-vis-derived ε_616_ below. The discrepancy between UV-vis and EPR estimates of ε_616_ systematically affects the magnitude of our estimated ΔE by 25 mV, but preserves the relative trend (EPR-based ΔE is reported in the **Supplementary Information Table S6)**. Incorporating NSQ quantitation, the E°_OX/NHQ_ values were determined as ‒318 mV for Q_103_K and ‒323 mV for Q_103_D at pH 7.9, shifts of +34 mV relative to the parental iLOV.

The difference in iLOV Q_103_X variants’ semiquinone accumulation indicates a change in the separation of E°_OX/NSQ_ and E°_NSQ/NHQ_ (ΔE). To determine the NSQ stability for iLOV and Q_103_X mutants, and hence ΔE, we performed comproportionation reactions, in which we mix equal concentrations OX- and NHQ-prepared iLOV and monitor the accumulation of NSQ via reaction **5** by UV-vis and EPR spectroscopy (**Figure 5** and **Supplemental Information Figure S13**). The comproportionation reactions were slow, reaching equilibrium in 10-20 h. Over the course of the reaction, we see a slight decrease in the OX state absorbance at 448 nm for iLOV, with a concomitantly small rise in the characteristic NSQ visible absorption feature at 616 nm (**Figure 5A**). The EPR spectrum of the iLOV OX/NHQ comproportionation equilibrium mixture showed an isotropic S = ½ singlet with a *g*_iso_ of 2.004 and a linewidth of 1.92 mT (**Figure 5B**). In general, ASQ radicals have a smaller linewidth of around 1.5 mT, while NSQ radicals have linewidths between 1.8 and 2.0 mT.^66^ Thus, the EPR and UV-vis spectra, and the pH dependence of the redox titrations, support an NSQ assignment. Compared to the Q_103_X mutants, the parental iLOV formed the least NSQ at equilibrium, suggesting that E°_OX/NSQ_ and E°_NSQ/NHQ_ are more crossed. Comproportionation reactions of Q_103_D produced the most NSQ (**Figure 5C**), with identical spectroscopic characteristics of the NSQ by EPR, *g*_iso_ of 2.004 and a linewidth of 1.94 mT (**Figure 5D**). To determine any contribution of ASQ to the comproportionation experiments, we measured the EPR spectrum of a Q_103_D comproportionation reaction at an even higher pH of 9.25, but observed no differences in either the EPR or UV-vis spectra (**Supplemental Information Figure S14**). These data imply significant differences in ΔE between iLOV and Q_103_D. The EPR spectra and simulation parameters for all comproportionation reactions are reported in the **Supplemental Information Figure S15** and **Table S5**.

**Figure 5.**
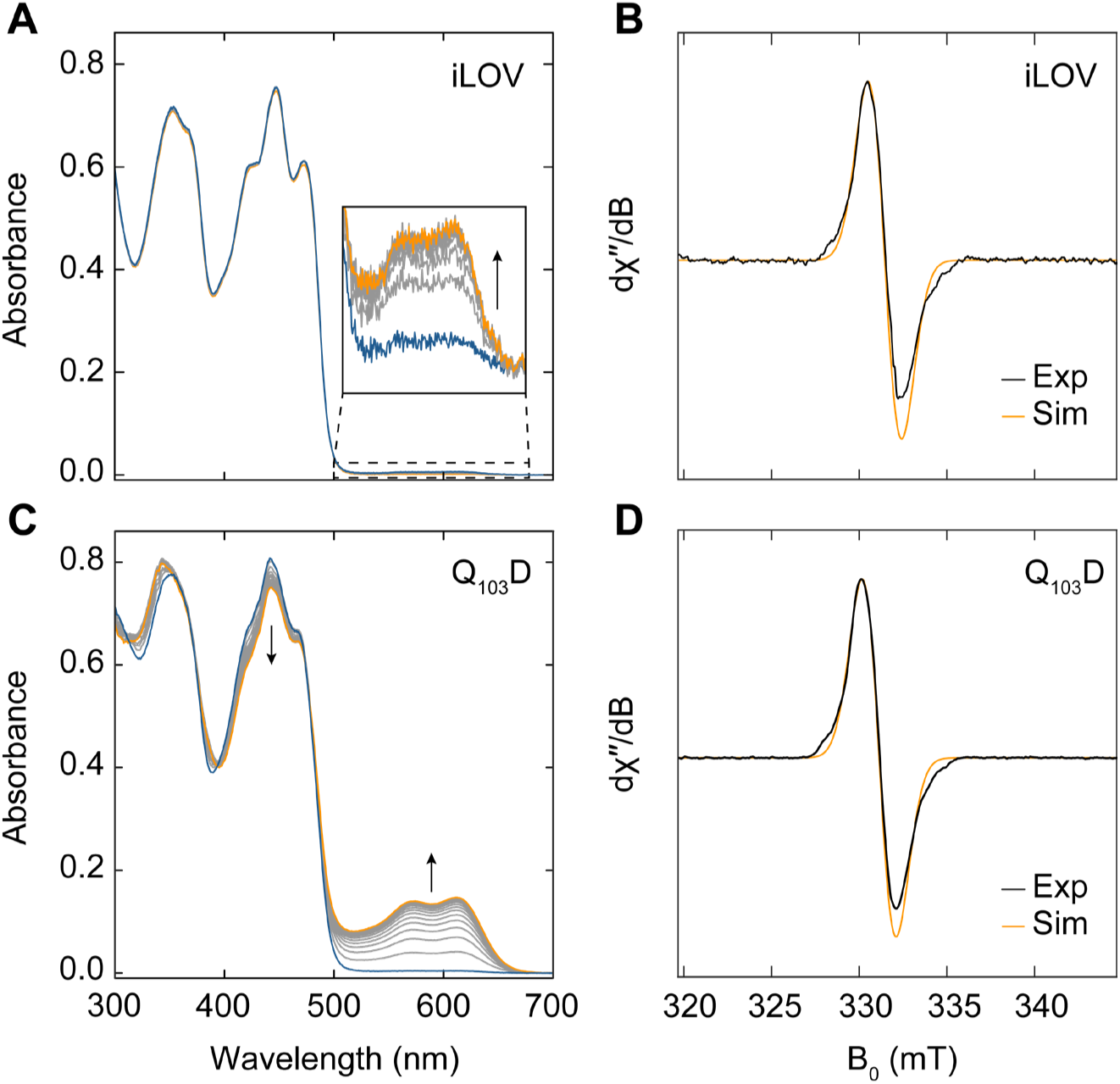
Comproportionation and characterization of the NSQ. **A** UV-vis time course of iLOV OX and NHQ comproportionation over 24 h. Arrows indicate the time-dependent direction of change in absorption from beginning (blue) to end (orange) of the reaction. **B** Normalized EPR spectrum (black) and simulation (orange) of the iLOV NSQ. **C** UV-vis time course of Q_103_D OX and NHQ comproportionation over 24 h. **D** Normalized EPR spectrum (black) and simulation (orange) of the Q_103_D NSQ.

With the measurement of iLOV and Q_103_X mutants’ E°_OX/NHQ_ in hand, and the relative *K*_eq_ of comproportionation, we calculated E°_OX/NSQ_ and E°_NSQ/NHQ_ across the series (**Table 2**). For iLOV, ΔE was ‒212 mV, corresponding to an E°_OX/NSQ_ of ‒458 mV and an E°_NSQ/NHQ_ of ‒246 mV. By comparison, Q_103_D exhibited a ΔE of ‒44 mV, corresponding to an E°_OX/NSQ_ of ‒342 mV and an E°_NSQ/NHQ_ of ‒298 mV. Thus, the Q_103_D mutation diminishes potential crossing relative to iLOV by increasing the E°_OX/NSQ_ by +116 mV, while also decreasing the E°_NSQ/NHQ_ by ‒52 mV. Across the mutants targeting N5, the E°_OX/NSQ_ was increased and the E°_NSQ/NHQ_ was decreased relative to iLOV, resulting in smaller ΔE. The shift in E°_OX/NSQ_ relative to iLOV contributed ≥ 70% to the overall change in ΔE, and the remaining 30% stemming from shifts to E°_NSQ/NHQ_, contributing 30% to the overall ΔE.

**Table 2.** ΔE, E°OX/NSQ, and E°NSQ/NHQ values for iLOV and Q103X mutants from spectrophotometric FMNOX and FMNNHQ comproportionation at pH 7.9. Propagated uncertainties in the last significant digit are reported in parentheses.

| iLOV variant | $E^{\circ}_{\text{OX/NHQ}}$ (mV) | $\Delta E$ (mV) | $E^{\circ}_{\text{OX/NSQ}}$ (mV) | % of $\Delta E_{\text{shift}}$ | $E^{\circ}_{\text{NSQ/NHQ}}$ (mV) | % of $\Delta E_{\text{shift}}$ |
| --- | --- | --- | --- | --- | --- | --- |
| iLOV | −354 | −212 (4) | −458 (2) | ref. | −246 (2) | ref. |
| Q<sub>103</sub>A | −337 | −176 (3) | −427 (2) | 86 | −251 (2) | 14 |
| Q<sub>103</sub>G | −328 | −112 (1) | −381 (1) | 85 | −259 (1) | 15 |
| Q<sub>103</sub>R | −321 | −127 (1) | −386 (1) | 77 | −269 (1) | 23 |
| Q<sub>103</sub>K | −318 | −104 (1) | −372 (1) | 79 | −269 (1) | 21 |
| Q<sub>103</sub>D | −323 | −44 (1) | −342 (1) | 69 | −298 (1) | 31 |
| FMN | −242<sup>a</sup> | −54 | −269<sup>a</sup> | N.A.<sup>b</sup> | −216<sup>a</sup> | N.A.<sup>b</sup> |

<sup>a</sup> Values from ref. 7.
<sup>b</sup> N.A. not applicable.

To provide insight into the molecular mechanism of flavin redox tuning by mutations proximal to the flavin N5, we performed classical MD simulations in both the OX and NSQ states. For the parental iLOV in the OX state, we ran multiple replicas of 500 ns trajectories. In most of those trajectories, the glutamine side-chain maintains a H-bonding interaction with the flavin O4′, with a representative replica (where H-bonding is maintained throughout most of the trajectory) giving a mean Nε•••O4′ distance of 3.2 Å and RMSF of 0.6 Å and mean Nε•••N5 distance of 4.0 Å and RMSF of 0.8 Å (see **Figure 6A** and **6B**).

**Figure 6.**
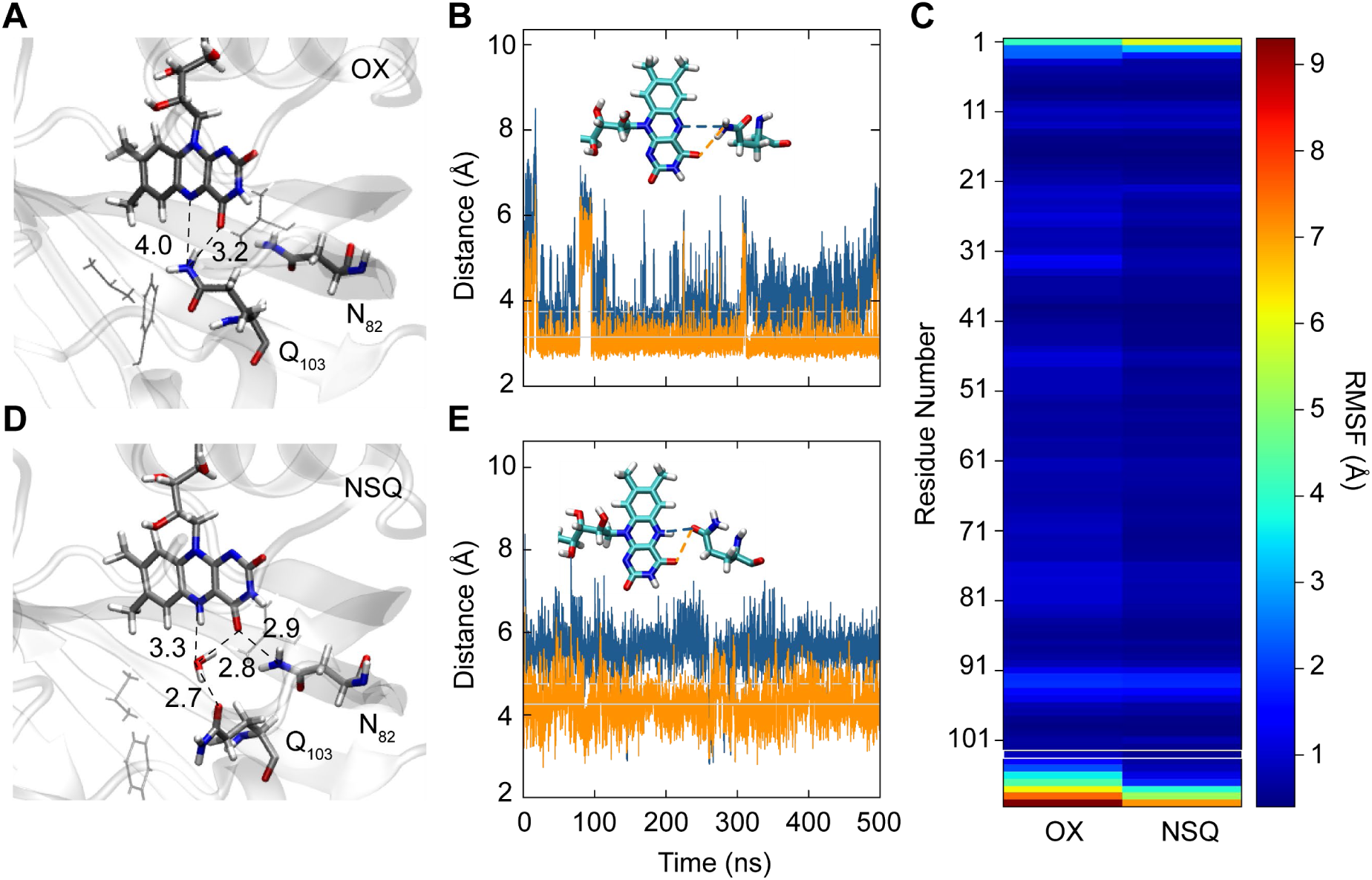
MD simulations of iLOV in the OX and NSQ state. **A** Representative structure of iLOV in the OX state from clustering analysis of a 500 ns MD trajectory. **B** Time-dependent distance analysis for Q_103_ amide N to the flavin N5 (blue) and O4′ (orange) of iLOV in the OX state from clustering analysis of a 500 ns MD trajectory. The solid gray line indicates the mean distance and the dashed line is the mean plus one RMSF. **C** Heat map showing the root-mean-square fluctuation (RMSF) of each residue in the OX and NSQ states of iLOV, with position 103 highlighted in a white box. **D** Representative structure of iLOV in the NSQ state from clustering analysis of a 500 ns MD trajectory. **E** Time-dependent distance analysis for Q_103_ amide carbonyl oxygen to the flavin N5 (blue) and O4′ (orange).

In some replicas of the iLOV OX state, we found that the glutamine side chain loses its H-bonding contact with the flavin O4′. This result is somewhat unsurprising in the context of the very hydrophobic environment of the Q_103_ side-chain in the X-ray structure, and the relative flexibility of this residue, as evidenced by the backbone Cα RMSD at this site near the C-terminus (**Figure 6C**).^24^ We note that in the full *A. thaliana* LOV2 domain of phototropin 2, from which iLOV is derived, Q_103_ is C-terminally flanked by the Jα helix, which may stabilize the hydrogen bonded state to N5/O4′ in the OX dark state, and the flipped out conformation resembles that which allosterically controls Jα helix unfolding in the light-activated state.^67,68^

We focus our analysis throughout the remainder of the manuscript on a replica where the H-bonding is maintained throughout the full 500 ns trajectory, consistent with the crystal structure (**Figure 3A**).

The reduction of iLOV to the NSQ state appears to result in less conformational flexibility of Q_103_, as indicated by the RMSF analysis in the C-terminal region (**Figure 6C**). Nevertheless, the conformation of Q_103_ does change in the NSQ state, rotating the side-chain such that its amide carbonyl oxygen, rather than the amide nitrogen, is pointed towards N5 (**Figure 6D**). This observation is consistent with what happens in native LOV domains in either the light-activated or reduced NSQ states.^68,69^ The flip comes at the expense of potential hydrogen bonding interactions. The lowest average distance for that of O4′•••O Q_103_ carbonyl is 4.3 Å with an RMSF of 0.5 Å and that for N5•••O is 5.6 Å with an RMSF of 0.5 Å (**Figure 6E**). In place of Q_103_ (strained) hydrogen bonds to the flavin N5/O4′, the calculations show that water enters the binding pocket in the NSQ state and mediates the H-bonding interactions between the flavin and Q_103_. Hydrogen bonding between N5 and a water molecule occurs with significantly higher probability (51.8%) in NSQ than the OX state (0.0%). The water occupancy within 4 Å from the flavin N5 in NSQ is significantly higher (94.4%) than the native OX state of iLOV (6.0%).

We next simulated the E°_OX/NSQ_ for the various Q_103_X mutants using alchemical relative free energy simulations, benchmarked against the parental iLOV. In previous work, a thermodynamic integration (TI) approach to simulating flavoprotein redox potentials yielded reduction potential errors on the order of < 1 kcal/mol (< 43 mV) for solvent-exposed residues in flavodoxin.^70,71^ While this approach is promising, the simulations here present added challenges that can contribute additional errors: (1) Q_103_ is directly hydrogen-bonded to the flavin and those hydrogen bonding interactions are expected to change upon N5 protonation, (2) some mutations involve large changes in the size and charge of the amino acid (*e.g.* protonated Q_103_R^(+)^), (3) Q_103_ is buried in a relatively hydrophobic environment, and (4) Q_103_ is near the flexible C-terminus. Such large perturbations at buried but dynamic residues present challenges to the requirement of BAR protocols where each λ step should weakly perturb the system.^72^ For these reasons, we opted to use longer MD simulations and 40 λ steps instead of the 8-14 λ steps used for flavodoxin.^48,70,73^

Our analysis of the alchemical trajectories and the resultant computed E°_OX/NSQ_ indicate that, even with a relatively long MD period and higher-density λ steps, errors between replicas were > 1 kcal/mol (**Supplemental Information Table S7** and **S8**). In the OX state, the free energy associated with introducing Q_103_X mutations yielded standard deviations (σ) between replicas on the order of 0.7-3.7 kcal/mol, with the greatest variations among replicas for the charged mutants Q_103_K^(+)^ (σ = 3.7 kcal/mol), Q_103_R^(+)^ (σ = 2.3 kcal/mol), and the deprotonated Q_103_D^(‒)^ (σ = 2.9 kcal/mol). Q_103_G also gave a larger error (σ = 2.1 kcal/mol) due to the large change in size of the amino acid. The Q_103_A and the Q_103_D^(0)^ mutations gave more reasonable errors (σ = 0.7 and 0.5 kcal/mol, respectively). In comparison to the OX state, σ between replicas of Q_103_X alchemical transformations in the NSQ state are smaller. For all NSQ mutations, σ ranged from 0.5 to 1.6 kcal/mol, with the exception of Q_103_K^(+)^, where σ was 3.5 kcal/mol. Nevertheless, since errors for the OX and NSQ states are additive when computing E°_OX/NSQ_ (giving σ of 1.9 kcal/mol to 7.2 kcal/mol, or 82 mV to 312 mV), these uncertainties are too large to reliably capture differences between the iLOV mutants.

Based on our observation with the parental iLOV, we suspected that solvent penetration dynamics may significantly contribute to the deviations among replicates. In all OX and NSQ simulations, the native iLOV binding pocket starts out anhydrous, but the gradual introduction of a mutation at Q_103_ causes water to enter the binding pocket near the flavin’s N5 region. **Figure 7A** shows the water cumulative number (CN), the number of waters within a variable radius shell centered at N5 at each λ step for one replica of Q_103_K^(+)^ NSQ state with a pre-equilibration time of 3 ns. Water penetration into the flavin binding site is indicated by an increase in CN at ≤ 4 Å. For Q_103_K^(+)^ specifically, we find different behavior between two replicas depending in the pre-equilibration time provided in the simulation (**Figure 7B**). In replica 1, which had a short pre-equilibration of only 3 ns at the λ = 0 step, water does not enter the binding pocket until later in the Q_103_→K transformation. As a result, water entering the binding pocket introduces a large perturbation and, after that point, the trajectories also show larger variations in the CN profile from one λ step to the other. These variations lead to large fluctuations in free energy not consistent with a gradually perturbative regime. By extending the pre-equilibration to 50 ns at the λ = 0 step this issue is mitigated; all λ trajectories behave more consistently after water enters the flavin binding pocket. A representative structure of the water-penetrated Q_103_K^(+)^ is shown in **Figure 7C**. After discarding the 3 ns trajectories for Q_103_K^(+)^, σ between replicas improved from 3.7 to 2.1 kcal/mol in the OX state, and from 3.5 to 1.0 kcal/mol in the NSQ state of Q_103_K^(+)^. We observed similar pre-equilibration time-dependent behavior for Q_103_R^(+)^ and consider only the longer pre-equilibration replicates further here for Q_103_R^(+)^ as well.

**Figure 7.**
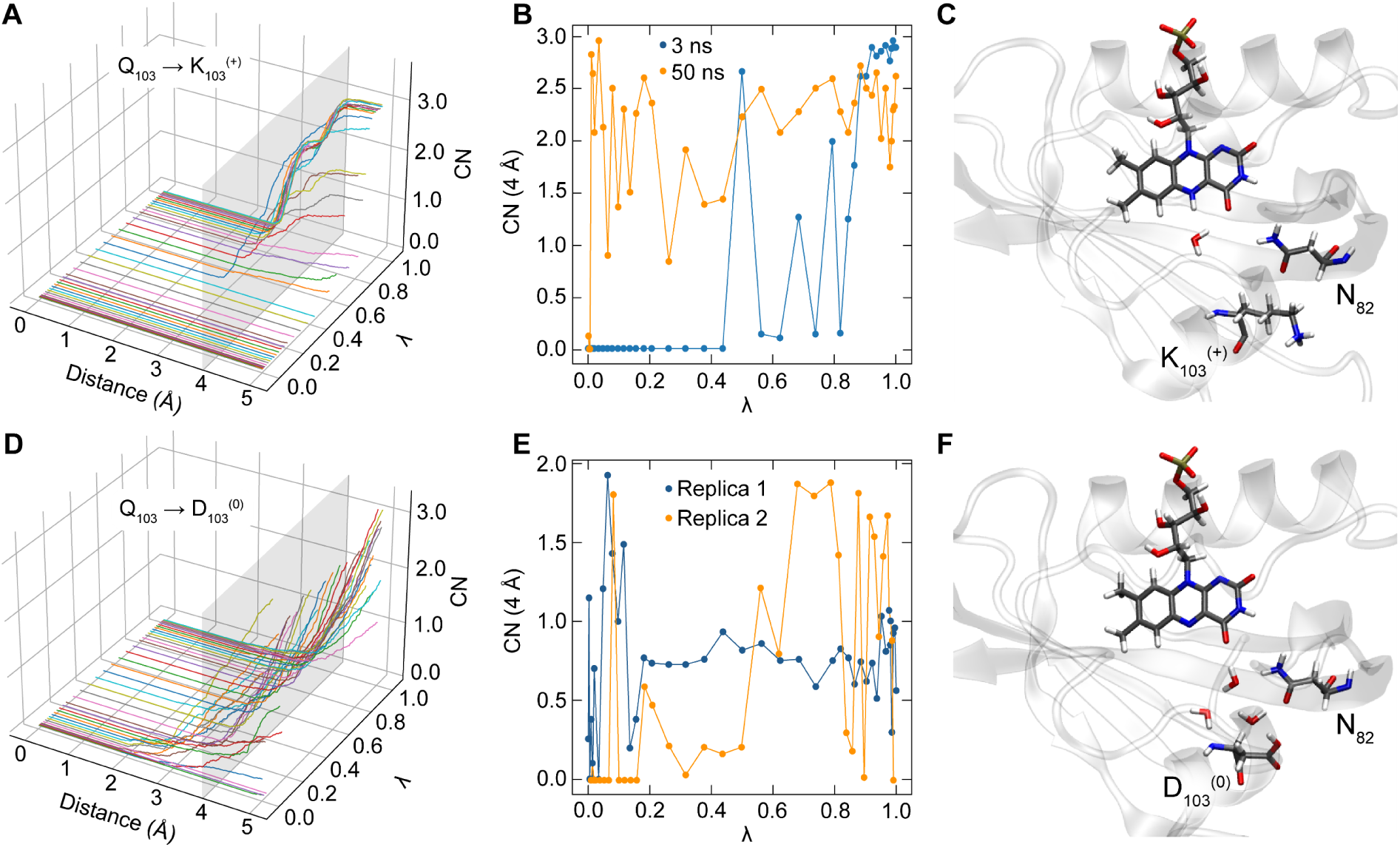
**A** Radial distribution of water cumulative number (CN) to N5 in the NSQ state during alchemical Q_103_ (λ = 0) to K_103_^(+)^ (λ = 1) transformation at each λ step with a 3 ns pre-equilibration time. **B** Comparison of CN at 4 Å during the Q_103_ to K_103_^(+)^ transformation for 3 ns (blue) and 50 ns (orange) pre-equilibration. **C** Representative structure of Q_103_K with water in the flavin pocket. **D** Radial distribution of water cumulative number (CN) to N5 during alchemical Q_103_ (λ = 0) to D_103_^(0)^ (λ = 1) transformation at each λ step with a 3 ns pre-equilibration time. **E** Comparison of CN at 4 Å during the Q_103_ to D_103_^(0)^ transformation for two different replicas with 50 ns pre-equilibration. **F** Representative structure of Q_103_D with water in the flavin pocket.

We also examined the solvent penetration behavior of the OX state of the protonated Q_103_D^(0)^ (**Figure 7D-7F**). Again, we consistently see water entering the binding pocket at some λ steps between 0→1. Both replicas display variations in the CN over the course of the alchemical transformation. In replica 1, water enters the binding pocket at an early λ step and remains inside the pocket until λ = 1. On the other hand, replica 2 displays intermittent entry of water into the flavin site, which leads to large fluctuations in free energy.

The examples of Q_103_K^(+)^ and Q_103_D^(0)^ are indicative of a larger theme, that a major source of error between different replicas of the same system is that not all replicas behave in the same way with regards to water entering the flavin binding pocket. However, we can qualitatively categorize the MD behavior into three empirical bins: (1) water enters the flavin binding pocket and remains there until the end in both the OX and NSQ state, (2) water initially enters the binding site at some intermediate λ step but is then ultimately expelled from the binding pocket at the end of the BAR protocol (at the final λ=1 step) in both the OX and NSQ states, and (3) water is expelled from the binding pocket in the OX state at the end of the BAR protocol but stays in the binding pocket in the case of NSQ. Categorizing replicas in this way, we notice that the comparison of computed E°_OX/NSQ_ from replicas belonging to each of those categories gives more internally consistent results. Specifically, using replicas that exclusively belong to category 3 gave considerably lower σ for almost all mutations. The only exceptions were Q_103_D^(0)^, where category 1 gave slightly lower error than category 3, and Q_103_R^(+)^, where not enough category 3 trajectories were obtained and both categories 1 and 3 are included.

Using the categorized replicas of each Q_103_X mutant and the thermodynamic cycle shown in **Figure 2** allowed us to calculate E°_OX/NSQ_ from 3-4 replicas (**Table 3**). The data from all replicas, including those not considered further herein, are reported in the **Supplemental Information Tables S7 and S8**. The calculated σ (sum of both OX and NSQ simulations) after binning are all ≤ 1.45 kcal/mol, instead of the 1.9 to 7.2 kcal/mol obtained using all replicas. Overall, the computed potentials are within one σ of the experimental values, the only exception being Q_103_K^(+)^, which is one of the two residues that introduces both a change in the charge state of the amino acid as well as a substantial increase in the side chain length. The general agreement of categories 1 and 3 with the experimental data and their internal consistency suggest that the flavin N5/O4′ in the NSQ state interacts with protein-penetrating water in all iLOV mutants, but to varying degrees. Indeed, there is good correlation between the positive potential shift of the E°_OX/NSQ_ couple and the water occupancy near N5/O4′ (**Table 3**). Furthermore, the protonated Q_103_D^(0)^ model does substantially better than the corresponding Q_103_D^(‒)^, with replicas yielding an E°_OX/NSQ_ closer to the experiment. These results indicate, in agreement with previous experimental and computational evidence,^33^ that the Q_103_D side-chain is protonated. Overall, the results suggest that the stability and frequency of hydrogen bonding to the NSQ state largely dictate the observed potential tuning of E°_OX/NSQ_.

**Table 3.** A comparison of simulated and experimental redox potentials. The simulated potentials are obtained using a thermodynamic cycle from shifts in redox potentials relative to iLOV by introducing mutations at the Q103 site. Standard deviations reported in parenthesis are the sum of σ from the OX and NSQ states.

| Variant | $\lambda$ Category | $E^{\circ}_{\text{OX/NSQ}}$ (mV) | | Difference (mV) | H<sub>2</sub>O Occupancy (N5, %) | H<sub>2</sub>O Occupancy (O4', %) |
| --- | --- | --- | --- | --- | --- | --- |
|  |  | Computed | Experimental |  |  |  |
| iLOV | N.A.<sup>a</sup> | ref.<sup>b</sup> | -458 | N.A.<sup>a</sup> | 6.0 | 7.5 |
| $Q_{103}A$ | 3 | -433 (31) | -427 | -6 | 1.4 | 9.3 |
| $Q_{103}G$ | 3 | -410 (63) | -381 | -29 | 3.4 | 36.4 |
| $Q_{103}R$ | 1, 3 | -391 (56) | -386 | -5 | 6.2 | 53.0 |
| $Q_{103}K$ | 3 | -444 (42) | -372 | -72 | 6.2 | 26.9 |
| $Q_{103}D^{(0)}$ | 1 | -364 (59) | -342 | -22 | 32.0 | 93.7 |
| $Q_{103}D^{(-)}$ | 3 | -442 (23) | | -100 | 10.8 | 81.0 |

<sup>a</sup> N.A. not applicable
<sup>b</sup> ref. reference value for integrated thermodynamic calculations

Representative structures of the mutated iLOV in the OX state, based on clustering analysis of the classical 500 ns trajectories, are shown in **Supplemental Information Figure S16**. The same figure also includes a heat map for the RMSF of individual amino acids. In addition to residue 103, we highlight two asparagines (N72 and N82) as well as one glutamine (Q44) that are involved in a hydrogen-bond network with the flavin and each other. In general, this hydrogen bond network is robust in most mutants, with Q44 and N72 in particular maintaining their conformational states across all mutants and clusters. However, N82, due to its proximity to residue 103, is more strongly affected by the mutations and can have different conformations. This is observed most notably in Q_103_R^(+)^ which, although is conformationally rigid, displays a different H-bonding pattern around N_82_ compared to other mutants.

The structures in the **Supplemental Information Figure S16** indicate that mutations at position 103 can introduce conformational changes distal to the N5 environment, either by allowing water into the binding pocket or by influencing flavin’s interactions with other neighboring amino acids. Such subtle (or in the case of N_82_, not so subtle) changes can propagate through the hydrogen bond network and affect hydrogen bonding at other flavin sites. Thus, the mutation of Q_103_ and N_82_ play the major role in forming the hydrogen bonding network near O4’/N5. Any mutation of Q_103_ perturbs the hydrogen bonding network such that either water or Q_103_X mutant has to form hydrogen bonding interactions with N_82_, otherwise N_82_ gets closer to the flavin and forms a hydrogen bond with O4′.

The heat map in the **Supplemental Information Figure S16** also shows that the mutants with the highest flexibility at the C-terminus are Q_103_D^(0)^ and Q_103_K^(+)^. Clustering analysis shows two major conformations of Q_103_D^(0)^. Representative structures from both conformations have a water molecule inside the active site, consistent with our earlier results indicating high water occupancy. On the other hand, for Q_103_K^(+)^, the two largest clusters are significantly different; one cluster has the lysine directly hydrogen bonded with flavin’s O4′, and the other cluster has the lysine side-chain pointing out away from the flavin and with a water molecule entering the binding pocket. The significant difference between those clusters can help explain why the redox potentials obtained from free energy simulations had larger standard deviations than for all other mutants.

## DISCUSSION

The ability of flavoproteins to tune the reduction potentials of their flavin cofactors to match those of their substrates or electron transfer partners underlies their remarkable catalytic versatility. Although flavin potentials are highly sensitive to the protein environment, the precise mechanisms by which flavoproteins modulate one- and two-electron couples remain unclear. Systematic investigations have been particularly challenging for crossed-potential flavins, where the inherent instability of the NSQ intermediate complicates measurement of individual one-electron potentials.

By targeting Q_103_ in iLOV, which resides near the flavin N5 and O4′ atoms, we demonstrated that single site mutations can significantly alter the degree of potential crossing (ΔE), spanning 164 mV across the Q_103_X series, while shifting the overall two-electron potential (E°_OX/NHQ_) by only ∼30 mV. The dominant contributor to these changes were positive shifts in the E°_OX/NSQ_ of up to +116 mV. This is somewhat unsurprising, given the proximity of the Q_103_X residue to the flavin N5, which becomes protonated upon reduction in the NSQ state. The trend implies that mutations to Q_103_ destabilize the OX state or stabilize the NSQ state, thereby raising E°_OX/NSQ_. Surprisingly, however, the effect of the Q_103_X mutations didn’t follow an immediately intuitive trend associated with the side-chain functional groups. Both acidic and basic residues showed similar E°_OX/NSQ_ (*e.g.* Q_103_K and Q_103_D), and elimination of hydrogen bonding entirely (*e.g.* Q_103_A and Q_103_G) had only modest effects on E°_OX/NSQ_ relative to the parental iLOV.

The E°_NSQ/NHQ_ couple was also affected by Q_103_X mutations, but to a lesser degree, with the most shifted couple in the Q_103_D mutant by ‒52 mV relative to the parent iLOV. In this, and all other Q_103_X cases, the E°_NSQ/NHQ_ couple becomes more negative, implying a stabilization of the NSQ and/or a destabilization of the NHQ state. Yet again, no clear correlation emerged with side-chain functional groups, underscoring the complexity of interpreting these changes without structural information across all redox states and detailed insights into conformational dynamics.

Our QM/MM and MD simulations help provide molecular context into the structural and dynamic differences between iLOV mutants, which is summarized in **Scheme 1**. For freely solvated FMN analog lumiflavin, hydrogen bonding to the flavin isoalloxazine ring heteroatoms is relatively unrestrained, and all atoms display some degree of hydrogen bonding in both the OX and NSQ state. Our numbers are similar to previous calculations with lumiflavin, with the exception of N1, which was not observed to form hydrogen bonds, likely due to the methyl group to N10 and the strong hydrogen bonds to O2′.^74^ In the X-ray crystallographic structure of iLOV with the FMN cofactor in the OX state, the hydrogen bonding interactions are significantly diminished in the hydrophobic flavin pocket.^24^ In particular no hydrogen bond < 3.0 Å is observed for either N1 or N5, destabilizing the cofactor. These interactions are largely reproduced in our extended MD simulations for Q_103_A, with the exception of hydrogen bonding interactions observed for N1, and the loss of the hydrogen bond to O4′ via loss of the Q_103_ side-chain. Overall, in the OX state, the isoalloxazine ring loses an average ‒1.0 (Δ_HB_) stabilizing hydrogen bond in moving into the iLOV binding pocket. The difference in hydrogen bonding between free and iLOV-bound FMN in the NSQ state are more pronounced. The hydrogen bonding to N5 and O4′ is substantially weakened, resulting in a net Δ_HB_ of ‒1.5. This greater destabilization of the NSQ partially explains the relatively low E°_OX/NSQ_ couple in iLOV relative to free FMN.

**Scheme 1.**
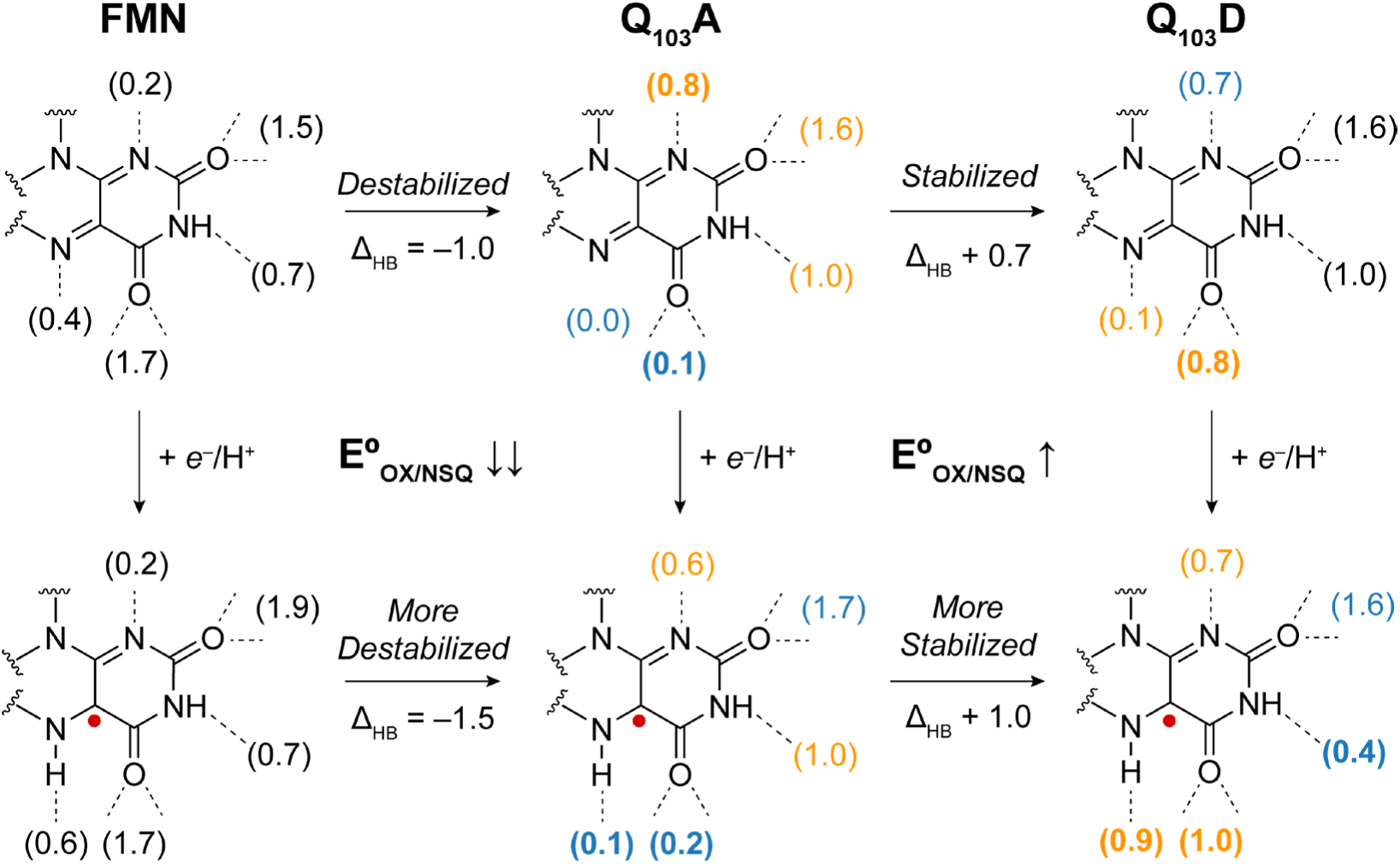
Proposed role of hydrogen bonding to the isoalloxazine ring affecting the FMN E°_OX/NSQ_ and E°_NSQ/NHQ_ for free FMN, Q_103_A, and Q_103_D. Average hydrogen bonds to the flavin heteroatoms from extended MD trajectories are shown in parentheses. Hydrogen bonding that increases (left to right) is indicated in orange and those that decrease in blue, with bold indicating a change > 0.5. Net change in hydrogen bonding (Δ_HB_) indicated for each transition.

Not all hydrogen bonds will contribute equally to the (de)stabilization of the FMN cofactor in iLOV. We expect the extent of stabilization of either OX or NSQ state through hydrogen bonding is dependent on the Δp*K*_a_ between donor and acceptor and steric constraints imposed by the protein scaffold.^75^ The p*K*_a_ of the FMN N5 in the OX state has been computationally estimated to be ‒ 10.^76^ When free in solution, hydrogen bonding between water and N5 is not particularly favored, with a Δp*K*_a_ ≥ 24,^77^ which is reflected in the low average hydrogen bond count of 0.4, but there are effectively no steric constraints. Conversely, the p*K*_a_ of the FMN NSQ is 8.6 and the associated Δp*K*_a_ for hydrogen bonds with a water acceptor is 8.6,^7,61^ indicating that hydrogen bonding to or from N5 in aqueous solution preferentially stabilizes the NSQ state over the OX state. We assume the p*K*_a_ of N5 in the OX state is not significantly perturbed within the flavin binding pocket (‒10), while the p*K*_a_ of the NSQ is clearly elevated to > 9, based on our EPR measurements of the Q_103_D SQ at high pH. As such, when comparing the change in hydrogen bonding between the free and iLOV-bound FMN, the loss of a hydrogen bond acceptor to N5-H in the NSQ state in iLOV is more destabilizing than the loss of a hydrogen bond donor to N5 in the OX state. The opposite argument has been made for the stabilizing effect of the NSQ state in flavodoxins, where a relatively strong hydrogen bond is formed between N5 and a glycine backbone amide carbonyl, increasing hydrogen bonding relative to free FMN.^78,79^

The FMN C=O4′ carbonyl bond is also weakened upon reduction to the NSQ state, making O4′ a better hydrogen bond acceptor.^80–84^ Even though hydrogen bonding interactions to O4′ are diminished in parental iLOV relative to free FMN in both the OX and NSQ state, we expect the destabilization effect to be greater for the NSQ state. Upon reduction, the aromaticity in the flavin ring loses some degree of delocalization, partially splitting along the N5-N10 axis. We expect that the localization of greater aromaticity to ring III will increase the impact of aromaticity-modulated hydrogen bonding at O4′, further destabilizing the NSQ relative to the OX state upon loss of H-bonding.^17^

On comprehensive comparison of the change in total hydrogen bonds (Δ_HB_, **Scheme 1**), the NSQ state is destabilized relative to the OX state by an average additional 0.5 hydrogen bonding interactions, notably at N5 and O4′, where relatively strong hydrogen bonds should form. The E°_OX/NSQ_ of iLOV is shifted ‒190 mV (18 kJ/mol) in iLOV relative to free FMN, which is consistent with a net loss of a hydrogen bond to the NSQ state. Additional effects of the protein binding pocket likely contribute to the E°_OX/NSQ_ shift as well, but those that affect both OX and NSQ states similarly will not result in shifts to E°_OX/NSQ_.^85–87^

Changes in hydrogen bonding character near the flavin N5 and O4′ across the Q_103_X mutations correlate with the E°_OX/NSQ_ shifts. For the Q_103_D mutation, which raise the E°_OX/NSQ_, hydrogen bonding increases at O4′, and to a lesser degree at N5, in the OX state, whereas hydrogen bonding increases significantly for both N5 and O4′ in the NSQ state. In both OX and NSQ, the effect of the Q_103_D mutation is to stabilize the FMN cofactor, but the NSQ is more stabilized by a ΔΔ_HB_ of 0.3. The shifts are relatively modest and due to the increased penetration of water into the flavin binding pocket proximal to N5/O4′. Water inside protein cavities is known to behave differently than that of the bulk, including its hydrogen bonding dynamics, p*K*_a_, and corresponding hydrogen bond energetics, and thus may not be directly comparable to solvation of N5 in free FMN.^85,88,89^ Nevertheless, by affecting both OX and NSQ oxidation states, their effect on the overall tuning of the E°_OX/NSQ_ couple (+116 mV, 11 kJ/mol) is more conservative than the net effect of moving FMN from bulk solvent to the iLOV pocket.

For the cationic mutants Q_103_R^(+)^ and Q_103_K^(+)^ we observe that the side-chain tends to flip out towards the solvent, which creates water penetration pathways in the NSQ state. The greater occupancy of water near N5 over the course of the simulations relative to Q_103_G correlate with the increase in E°_OX/NSQ_. While electrostatics can tune flavin redox potentials, these effects disproportionately affect proton-uncoupled redox transformations in which an ASQ or AHQ is formed.^20,90^ Our data here are consistent with this, in that, the introduction of an expectedly charged residue does little to change the reduction potential of an charge-neutral OX→NSQ or NSQ→NHQ transition. Across the series of Q_103_R^(+)^, Q_103_K^(+)^, and Q_103_D^(0)^, the MD simulations indicate water occupancy and hydrogen bonding correlate well with the computed E°_OX/NSQ_, providing a rationale and experimentally testable model for flavin redox tuning via hydrogen bonding interactions to N5 and O4′.

For Q_103_D, only the neutral state was found to agree with experiment, but with one methylene carbon fewer than the Q side-chain, no hydrogen bond can be formed with N5. Instead, water overwhelmingly penetrates into the flavin binding pocket, where it can hydrogen bond with N5 and O4′, and is additionally stabilized by hydrogen bonding interactions with the carboxylic acid side-chain carbonyl of Q_103_D^(0)^. This hydrogen bonding interaction likely polarizes the water molecule, making it a better hydrogen bond acceptor within the hydrogen bonding network.

The iLOV scaffold and Q_103_X mutations also change the bound FMN E°_NSQ/NHQ_, relative to the free cofactor. In the parental iLOV, E°_NSQ/NHQ_ is only 30 mV below that of free FMN. The p*K*_a_ of the free riboflavin N1 in the NSQ state is 2.3,^91^ and shifts to 6.7 in the NHQ state, corresponding to a Δp*K*_a_ in H_2_O•••H-N1 hydrogen bonds of 11.7 and 6.7, respectively. This Δp*K*_a_ difference between the two oxidation states is more modest than for N5 between OX and NSQ states, indicating that hydrogen bonding contributes to stabilization of both NSQ and NHQ states, but more so in the NHQ. The X-ray structure of iLOV (in the OX state) contains a strained hydrogen bond between the amide -NH_2_ of Q_44_ and N1 (N•••N distance of 3.4 Å) and a shorter hydrogen bond with O2′ (N•••O distance of 2.9 Å), analogous to the Q_103_ interactions with N5 and O4′, whereas our MD simulations show the Q_44_ amide -NH_2_ forms hydrogen bonds to N1 in about 60-80% of frames, depending on oxidation state. This is in contrast to the situation in water, where only 20% of frames form hydrogen bonds at N1. Assuming Q_44_ can act as both a hydrogen bond donor and acceptor during the NSQ→NHQ transition, hydrogen bonding capacity to N1 is maintained in both NSQ (acceptor) and NHQ (donor) states. Here, we predict that hydrogen bonding will stabilize both NSQ or NHQ similarly, explaining the small change in E°_NSQ/NHQ_ relative to free FMN. Nevertheless, there are other factors the protein environment likely imposes on the NSQ→NHQ transition that are distinct from those experienced in bulk water, such as the flavin “butterfly” bend in the NHQ state, which could partially explain the slightly more negative E°_NSQ/NHQ_, but is not addressed in the experiments or simulations presented herein.

The trend in E°_OX/NSQ_ across the Q_103_X mutants was also reflected in the E°_NSQ/NHQ_, but instead decreases the reduction potential from iLOV to Q_103_D, indicating that the mutations either stabilize the NSQ or destabilize the NHQ. The distal nature of the Q_103_X mutations relative to N1 makes changes in hydrogen bonding to N1 across the series unlikely. We hesitate to speculate on the molecular origin of this trend, but note that the p*K*_a_ of N5 in the NHQ state has never been measured to our knowledge, but is likely extremely high (> 14), making any N5 hydrogen bonds minimal in the overall NHQ stabilization. In this scenario, systematic increases in stability of the NSQ due to hydrogen bonding to water, as we argue for the tuning of E°_OX/NSQ_, should decrease E°_NSQ/NHQ_, assuming little change in the stability of the NHQ state across the series. This hypothesis will be tested in future studies.

It is worthwhile to discuss the methodological approach to the alchemical free energy calculations more thoroughly, as these methods may be useful in other redox calculations in complex and dynamic redox proteins and enzymes. First, we notice that the computed reduction potentials reported for all mutants in **Table 3** are systematically more negative than the experimentally measured potentials. One possible explanation for such a systematic error is the different force fields for the OX and NSQ flavin states, even though the charges and geometries were obtained from QM/MM calculations. Another source of error may be introduced by the difference in solvation of those two states, since in most simulations (*e.g.* belonging to category 3) the OX of the final mutated protein at λ = 1 is dry while for NSQ it is solvated. As more information is collected on flavin redox tuning, these types of experiments can serve as an experimental benchmark from which to compare computational predictions of the reduction potentials of flavoproteins. While several free energy studies have been reported for computing redox potentials or effects of mutations on protein interface (in flavoproteins and other related biological systems), these protocols have often relied on shorter dynamics and fewer λ steps.^48,70,73^ Such protocols can be suitable for systems like flavodoxin, where flavin is already solvent-exposed. Here, even with 40 λ steps (inclusion of more intermediate states connecting the end points) and longer dynamics (600 ns per replica) than is typically used, we find larger errors in the calculations. This is because free energy protocols are suitable when a change is introduced smoothly and gradually. When flavin is engulfed by a protein environment, the alchemical transformation is more easily disrupted by random and intermittent entering/exiting of water near the flavin. Even with these errors, it is possible to obtain redox potentials in biomolecules with errors on the order of 1-2 kcal/mol, but this requires extensive sampling, multiple replicas to identify outliers, and careful analysis of the trajectories to monitor issues like water access and other indicators of large perturbations occurring along the λ steps. Often, computing redox potentials in solution can be done with reasonably high accuracy using implicit solvation,^92^ but here we instead highlight both the challenges, as well as the capability to compute redox potential in a complex and heterogeneous environment such as iLOV. The experiments reported here, which studies the effect of a series of single-point mutations on the reduction potential of iLOV, can serve as a benchmark set for computational methods being developed such as QM/MM MD simulations,^93^ polarizable force fields,^94^ and machine learning approaches.^95,96^

## CONCLUSION

In this study, we tuned the reduction potential of a flavin cofactor embedded within the small fluorescent flavoprotein iLOV by changing the identity of a side chain residue nearby the N5 and O4′ atoms of the isoalloxazine ring. We showed that the identity of this side chain and the corresponding increase in solvent penetration to the N5 pocket reduced the extent of potential crossing of iLOV, while only modestly affecting the overall two-electron potential of the cofactor. This can help serve as a reference from which to gauge the accuracy of quantum mechanical predictions of flavoprotein reduction potentials and can better inform future engineering of artificial flavoproteins via the thermodynamics of proton-coupled electron transfer.

## ASSOCIATED CONTENT

### Supporting Information

Initial and post-reconstitution FMN/protein content, protein purification and purity gels, iLOV Q_103_X mutagenic primers, photoreduction and ε_616_ determination for the Q_103_D NSQ, statistical clustering analysis of MD trajectory, ^1^H NMR spectra and assignment of safranine isomers, cyclic voltammograms and Pourbaix diagram of safranine isomer 3, excitation spectra of iLOV and Q_103_X mutants, emission lifetimes for iLOV and Q_103_X mutants, emission lifetime fit constants and quantum yields, protein film voltammetry of iLOV and Q_103_X mutants, xanthine/xanthine oxidase redox titrations of iLOV and Q_103_X mutants, equilibrium redox titrations of all Q_103_X mutants, X-band EPR quantitation of Q_103_D NSQ, comproportionation approach to equilibrium for iLOV and Q_103_X mutants, X-band EPR spectra of iLOV and Q_103_X mutants NSQ, X-band EPR spectrum of Q_103_D NSQ at pH 9.25, EPR simulation parameters for iLOV and Q_103_X mutant NSQs, and comparison of ΔE estimates using UV-vis vs. EPR NSQ quantitation, free energies of mutation for OX and NSQ states of iLOV and Q_103_X, computed reduction potentials for Q_103_X mutations, and Statistical clustering analysis and representative structures of Q_103_X

The following files are available free of charge.

## AUTHOR INFORMATION

### Author Contributions

The manuscript was written through contributions of all authors. All authors have given approval to the final version of the manuscript.

## ACKNOWLEDGMENT

We gratefully acknowledge Collin Origer for help in preparing iLOV proteins and Alexander Mikhailovsky for emission lifetime measurements. We are also grateful to Jacopo D’Ascenzi for aid in developing and early testing of the free energy protocol used in this work. We also thank Prof. Massimo Olivucci for useful discussions on the computational methodology. This material is based upon work supported by the National Science Foundation (NSF) under Grant CHE-2047667 (S.G.). S.E. and S.G. acknowledge use of Advanced Research Computing Technology and Innovation Core (ARCTIC) resources at Georgia State University’s Research Solutions, made available by the NSF Major Research Instrumentation (MRI) grant number CNS-1920024.

---

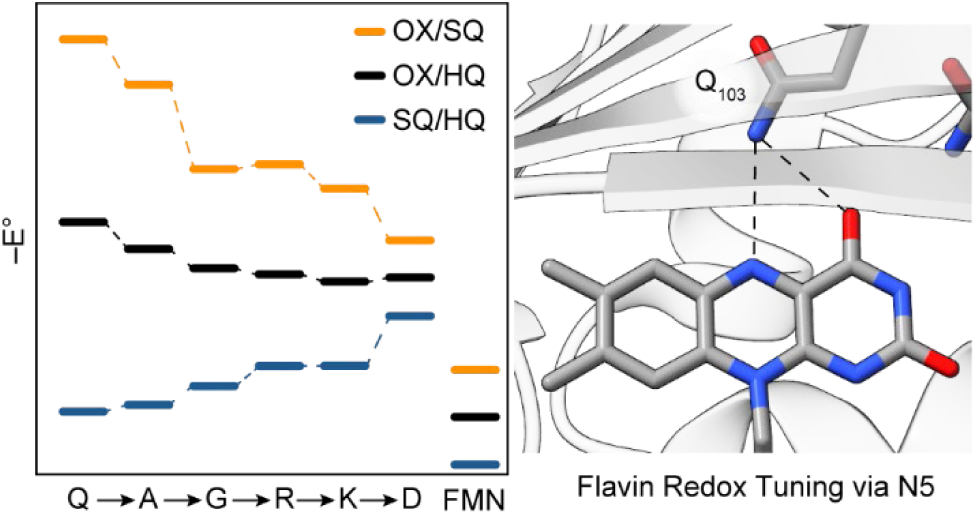

