## Supplemental Information for "Measurement and Control of Crossed Potentials in a Flavoprotein"

| Contents | Page |
|---|---|
| **Table S1.** Initial and post-reconstitution FMN/protein content | 3 |
| **Figure S1.** Protein purification and purity gels | 4 |
| **Table S2.** iLOV $Q_{103}X$ mutagenic primers | 5 |
| **Figure S2.** Photoreduction and $\varepsilon_{616}$ determination for the $Q_{103}D$ NSQ | 6 |
| **Figure S3.** Statistical clustering analysis of MD trajectory frames | 7 |
| **Figure S4.** $^{1}H$ NMR spectra of safranine isomers | 8 |
| **Table S3.** $^{1}H$ NMR spectral assignments of safranine isomers | 9 |
| **Figure S5.** Cyclic voltammograms and Pourbaix diagram of safranine isomer 3 | 10 |
| **Figure S6.** Excitation spectra of iLOV and $Q_{103}X$ mutants | 11 |
| **Figure S7.** Emission lifetimes for iLOV and $Q_{103}X$ mutants | 12 |
| **Table S4.** Emission lifetime fit constants and quantum yields | 13 |
| **Figure S8.** Protein film voltammetry of iLOV and $Q_{103}X$ mutants | 14 |
| **Figure S9.** Xanthine/xanthine oxidase redox titrations of iLOV and $Q_{103}X$ mutants | 15 |
| **Figure S10.** Equilibrium redox titrations of all $Q_{103}X$ mutants | 16 |
| **Figure S11.** Oxidative titration of iLOV with Ferricyanide | 17 |
| **Figure S12.** X-band EPR quantitation of $Q_{103}D$ NSQ | 18 |
| **Figure S13.** Comproportionation approach to equilibrium for iLOV and $Q_{103}X$ mutants | 19 |
| **Figure S14.** X-band EPR spectrum of $Q_{103}D$ NSQ at pH 9.25 | 20 |
| **Figure S15.** X-band EPR spectra of iLOV and $Q_{103}X$ mutants NSQ | 21 |
| **Table S5.** EPR simulation parameters for iLOV and $Q_{103}X$ mutant NSQs | 22 |
| **Table S6.** Comparison of ΔE estimates using UV-vis vs. EPR NSQ quantitation | 23 |
| **Table S7.** Free energies of mutation for OX and NSQ states of iLOV and $Q_{103}X$ | 24 |
| **Table S8.** Computed reduction potentials for $Q_{103}X$ mutations | 26 |
| **Figure S16.** Statistical clustering analysis and representative structures of $Q_{103}X$ | 28 |
| **References.** | 29 |

**Table S1**. As-purified and reconstituted FMN loading of iLOV variants. FMN concentrations were determined via UV-vis absorption at 448 nm using an extinction coefficient of 14,800 $M^{-1}$ $cm^{-1}$. For blue-shifted mutants ($Q_{103}$K/D/R/A/G) the absorption at 440 nm using an extinction coefficient of 16,100 $M^{-1}$ $cm^{-1}$ was used. Protein concentration was determined via the Pierce 660 assay. Initial FMN:iLOV ratios prior to flavin reconstitution are based on single measurements. Reconstituted FMN:iLOV ratios and uncertainty are based on the average of triplicate measurements and error is reported as the standard deviation.

| iLOV Variant | As-purified FMN:iLOV | Reconstituted FMN:iLOV |
|---|---|---|
| WT | 0.21 | 1.05 ± 0.02 |
| $Q_{103}$K | 0.27 | 0.94 ± 0.02 |
| $Q_{103}$D | 0.36 | 1.14 ± 0.02 |
| $Q_{103}$R | 0.31 | 0.96 ± 0.02 |
| $Q_{103}$G | 0.44 | 1.09 ± 0.01 |
| $Q_{103}$A | 0.27 | 1.12 ± 0.02 |

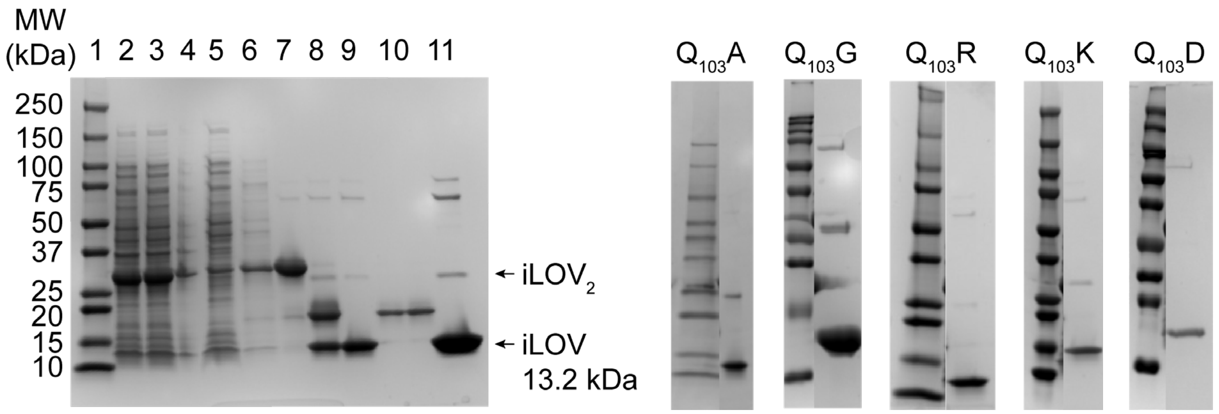


**Figure S1**. Purification (right) and purity (left) SDS-PAGE gels of iLOV and $Q_{103}X$ variants. Left: Typical purification gel for iLOV. Lanes as follows: (1) Precision Plus Molecular Weight Standard, (2) total cell lysate, (3) soluble fraction of cell lysate, (4) insoluble pellet of cell debris, (5) Ni-NTA flow through (pre-ULP1 digestion), (6) Ni-NTA wash, (7) Ni-NTA elution ($(H)_6$,-SUMO-iLOV), (8) ULP1 digestion (iLOV and $(H)_6$-SUMO), (9) digested Ni-NTA flow through and wash (iLOV), (10) ULP1 digestion Ni-NTA elution ($(H)_6$-SUMO), run in duplicate, and (11) buffer exchanged purified iLOV. Heavier molecular weight bands around 26 kDa, but slightly lower than $(H)_6$,-SUMO-iLOV, present in the purified fraction are small quantities of photo-oxidized and covalently crosslinked iLOV dimers.[1]

**Table S2**. Site directed mutagenesis (SDM) primer sequences for iLOV variants.

| iLOV Variant | Primer Sequences |
|---|---|
| $Q_{103}K$ | (forward) 5′-gcagtatttcatcggtgttaagctggatggttctga-3′<br>(reverse) 5′-ccagcttaacaccgatgaaatactgc-3′ |
| $Q_{103}D$ | (forward) 5′-ggtgttgatctggatggttctgaccacg-3′<br>(reverse) 5′-gaaccatccagctgaacatcgatgaaatactg-3′ |
| $Q_{103}R$ | (forward) 5′-gcagtatttcatcggtgttcggctggatg-3′<br>(reverse) 5′-gaaccatccagccgaacaccgatg-3′ |
| $Q_{103}G$ | (forward) 5′-cggtgttgggctggatggttctgaccacg-3′<br>(reverse) 5′-cagcccaacaccgatgaaatactgcagctcgcc-3′ |
| $Q_{103}A$ | (forward) 5′-cggtgttgcgctggatggttctg-3′<br>(reverse) 5′-gaaccatccagcgcaacaccgatg-3′ |

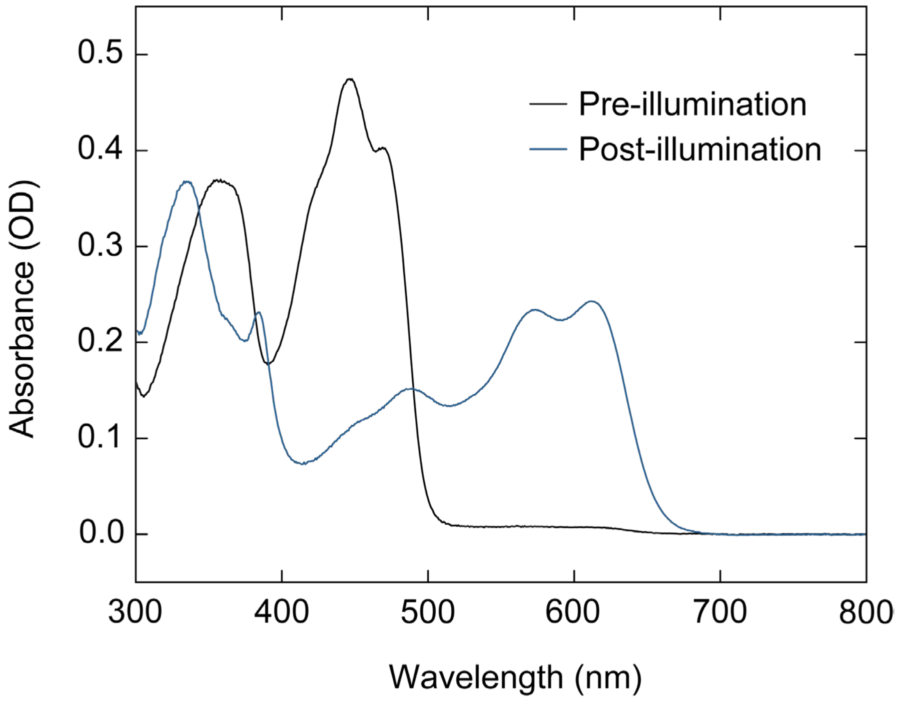


**Figure S2**. Determination of the $Q_{103}D$ NSQ extinction coefficient at 616 nm. Samples of $Q_{103}D$, initially in the $FMN_{OX}$ state (black), were illuminated with 450 nm light and photo-reduced until the absorbance at 616 nm reached a maximum (blue). The ratio of the absorbance between 616 and 444 nm was multiplied by the extinction coefficient at 444 nm (16,100 ± 200 $M^{-1}$ $cm^{-1}$).

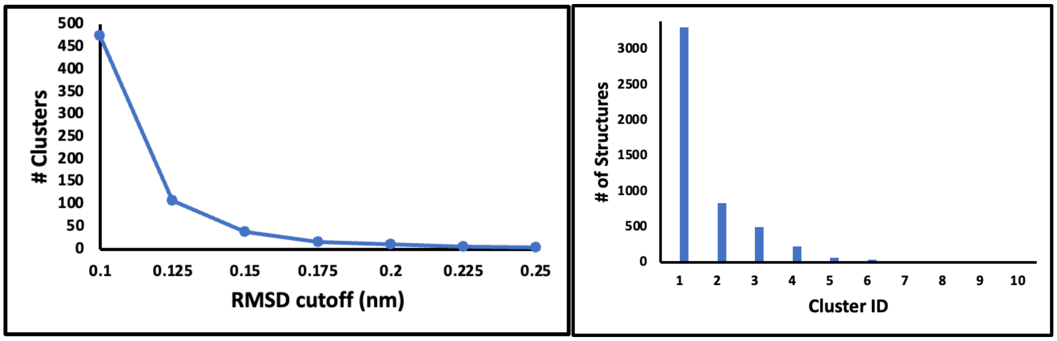


**Figure S3**. Statistical clustering analysis based on RMSD cutoff (left), and number of structures per cluster with an RMSD cutoff of 0.2 nm (right). Ten total clusters were obtained and only 2 had more than 500 MD snapshots out of the total 5001 frames.

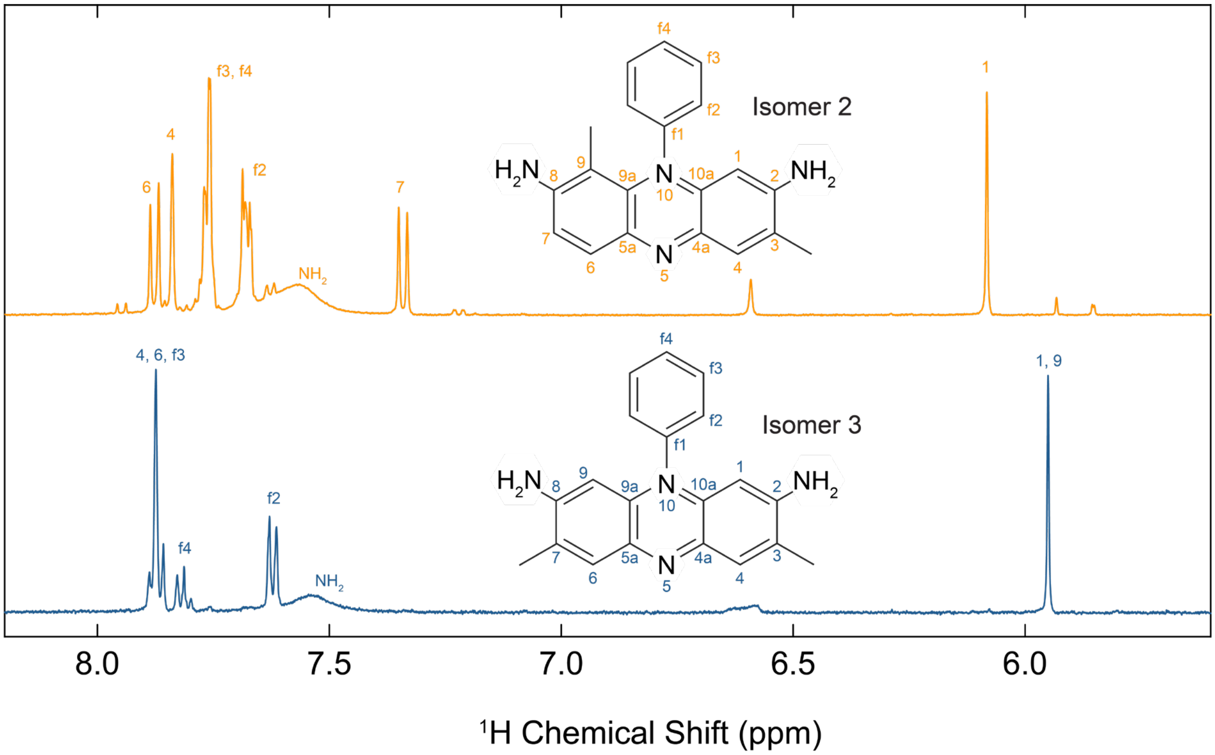


**Figure S4**. $^1$H NMR spectra and resonance assignments of safranine isomers in $d_6$-DMSO in the aromatic region. Unassigned peaks in the Isomer 2 spectrum are thought to be residual contaminants that were not separated during HPLC-MS purification.

**Table S3**. One-dimensional $^{1}H$ NMR spectral assignments of safranine isomers.

| |
|---|
| **Isomer 2** |
| $^{1}$H-NMR ($d_6$-DMSO, 500 MHz): δ 1.34 [3H, s, C(9)C$H_3$], 2.29 [3H, s, C(3)C$H_3$], 6.08 [1H, s, *H*(1)], 7.34 [1H, d, J = 9.2 Hz, *H*(7)], 7.68 [2H, m, *H*(f2)], 7.77 [3H, m, *H*(f3) & *H*(f4)], 7.84 [1H, s, *H*(4)], 7.88 [1H, d, J = 9.2 Hz, *H*(6)]. |
| **Isomer 3** |
| $^{1}$H-NMR ($d_6$-DMSO, 500 MHz): δ 2.30 [6H, s, C(3/7)$CH_3$:(*H*(17))], 5.95 [2H, s, *H*(1/9)], 7.62 [2H, d, J = 7.5 Hz, *H*(f2)], 7.81 [1H, t, J = 7.4 Hz, *H*(f4)], 7.87 [4H, m, *H*(4/6) & *H*(f3)]. |

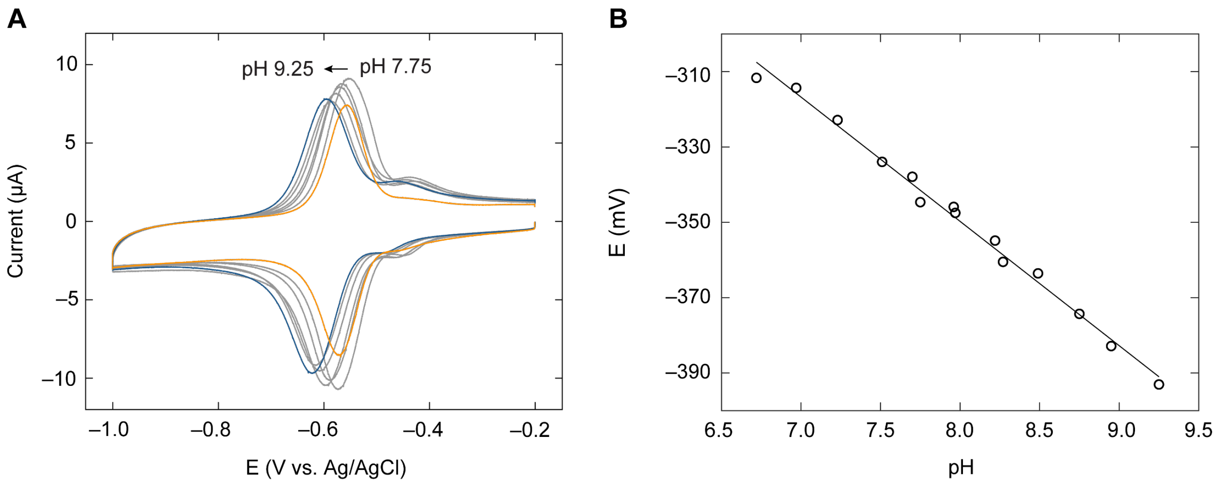


**Figure S5**. Pourbaix analysis of safranine isomer 3. **A** Representative cyclic voltammetry (CV) scans of safranine isomer 3 from pH 7.75 (orange) to 9.25 (blue). **B** Pourbaix diagram (midpoint potential vs. pH) of safranine isomer 3 and associated linear fit. The slope of the fit was –33 mV/pH (1 $H^+$/2 $e^-$) with an $R^2$ of 0.99.

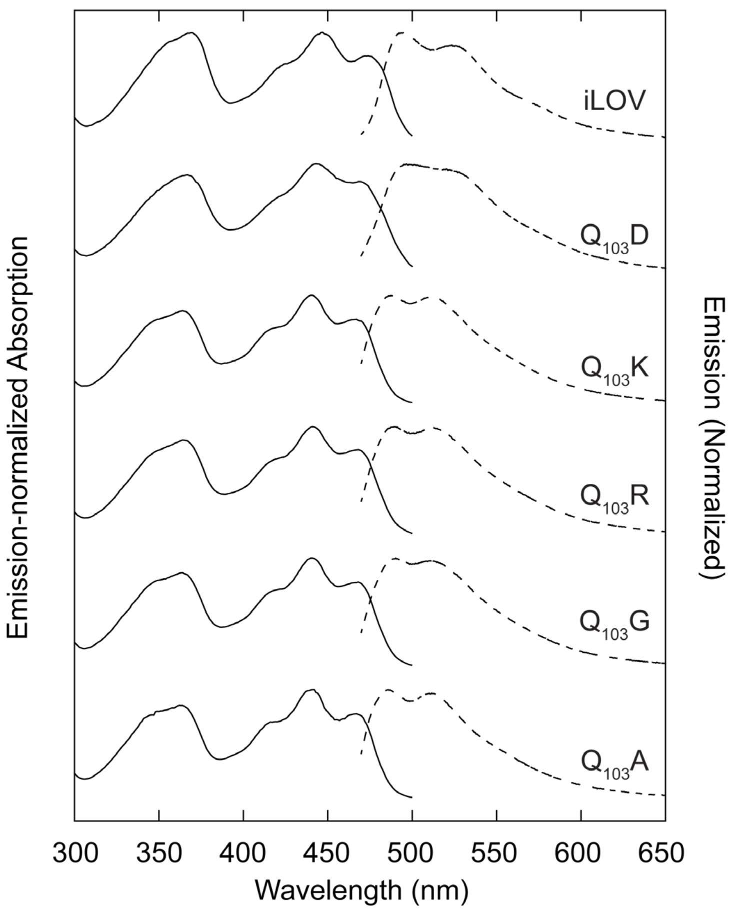


**Figure S6**. Emission-normalized ($\lambda_{em}$ = 520 nm) absorption spectra (excitation spectra, solid lines) of iLOV and $Q_{103}X$ mutants. Emission spectra ($\lambda_{ex}$ = 450 nm, dashed lines) are the same as those reported in the main text Figure 2.

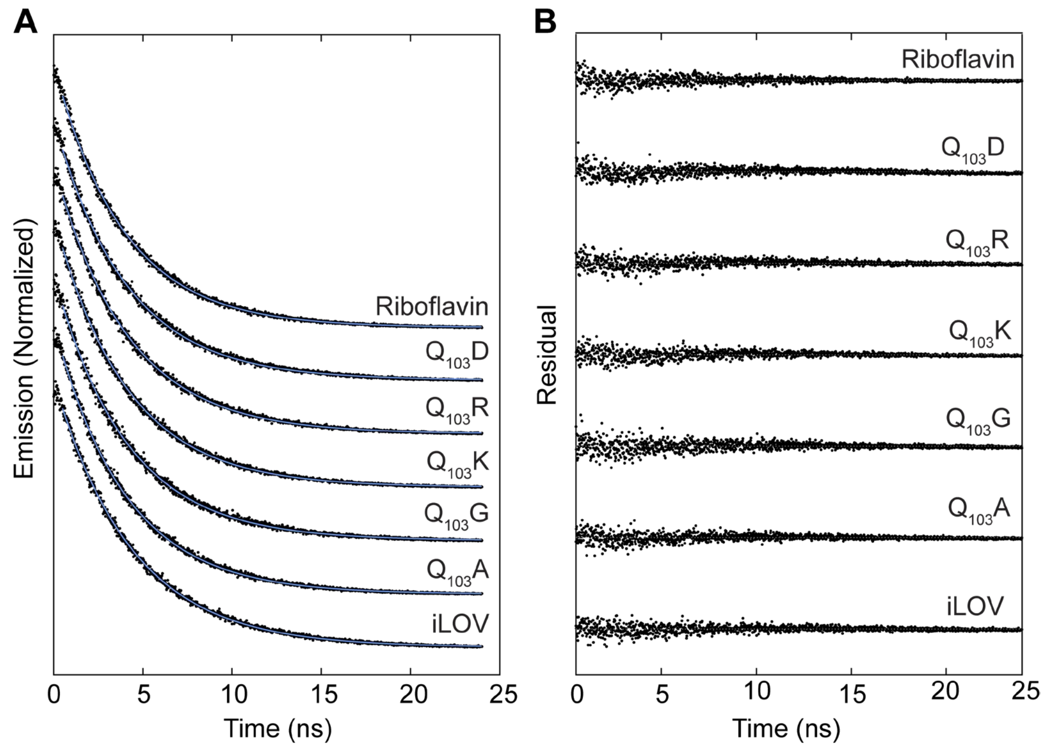


**Figure S7**. Fluorescent lifetimes of iLOV, $Q_{103}$X mutants, and riboflavin. **A** Emission lifetime traces (dots, $\lambda_{ex}$ = 400 nm) and associated single exponential fits to the experimental data (blue line). **B** Residuals of the experimental data relative to the fit.

**Table S4.** Fluorescence decay lifetimes (τ) and quantum yields (Φ) and emission maxima ($\lambda_{max}$) of iLOV and $Q_{103}X$ variants.

| iLOV Variant | τ (ns) | Φ | $\lambda_{max}$ |
|---|---|---|---|
| iLOV | 4.37 (0.01)[1] | 0.37 (0.01)[2] | 492 |
| $Q_{103}A$ | 4.01 (0.01) | 0.37 (0.01) | 486 |
| $Q_{103}G$ | 4.09 (0.01) | 0.37 (0.01) | 490 |
| $Q_{103}K$ | 4.07 (0.01) | 0.35 (0.01) | 488 |
| $Q_{103}R$ | 4.11 (0.01) | 0.36 (0.01) | 489 |
| $Q_{103}D$ | 4.05 (0.01) | 0.33 (0.01) | 498 |
| Riboflavin | 3.92 (0.01) | 0.26[3] | |

[1] Numbers in parentheses represent the standard deviation resulting from a fit of a single exponential decay trace.
[2] Numbers in parentheses represent the error propagation of standard deviation resulting from the integration of triplicate measurements of the iLOV variant emission spectra and the standard deviation resulting from their respective absorbance measurements.
[3] Values taken from Ref. 2.

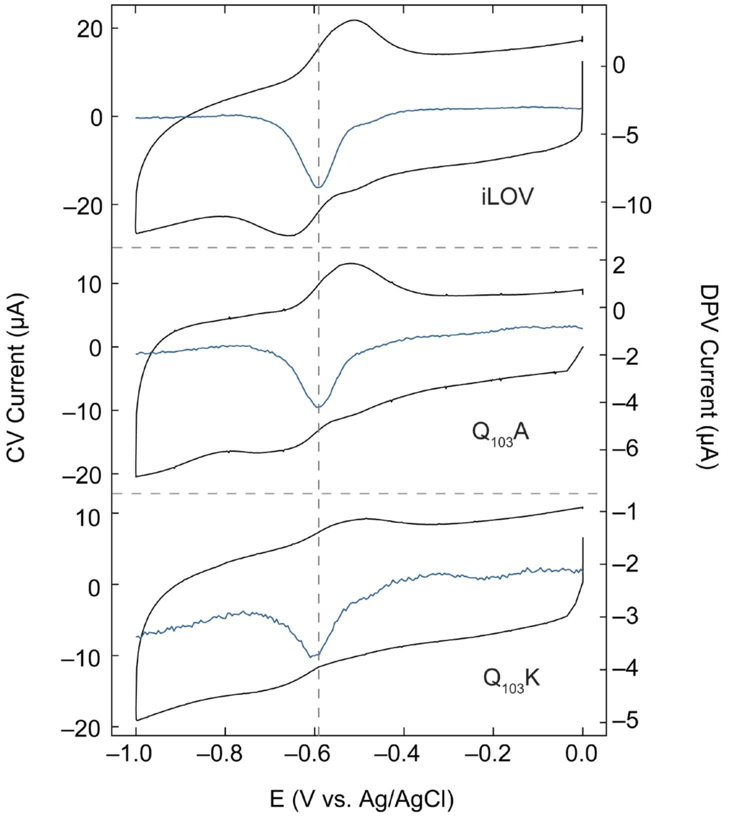


**Figure S8**. Initial attempts to measure $E°_{OX/NHQ}$ for iLOV variants using protein film voltammetry. Cyclic voltammograms (CV, black) and differential pulse voltammograms (DPV, blue) are overlayed. The voltametric setup was identical to that of the safranine measurements. About 5 µL of 1 mM protein was cast onto the surface of a pyrolytic graphite edge-plane working electrode and allowed to dry over a flow of nitrogen. After 1 hour of drying, the electrode was immersed in an electrode solution that was degassed by sparging with Argon gas for 30 minutes prior. A Ag/AgCl (3M NaCl) reference electrode and a platinum wire counter electrode were used. CVs were scanned from 0 to –1.0 V with a scan speed of 1 V/s while DPVs were collected over the same range with a pulse height of 25 mV, a pulse width of 10 ms, a step height of –5 mV and a step time of 100 ms. CVs contained several odd features. The cathodic peak was lower in intensity and broader than the anodic peak, both peak widths were unusually broad at ~150 mV, and all traces contained an unusual peak-to-peak separation of around 140 mV (as opposed to the expected value of 0 mV for an adsorbed analyte). No differences were observed between mutants.

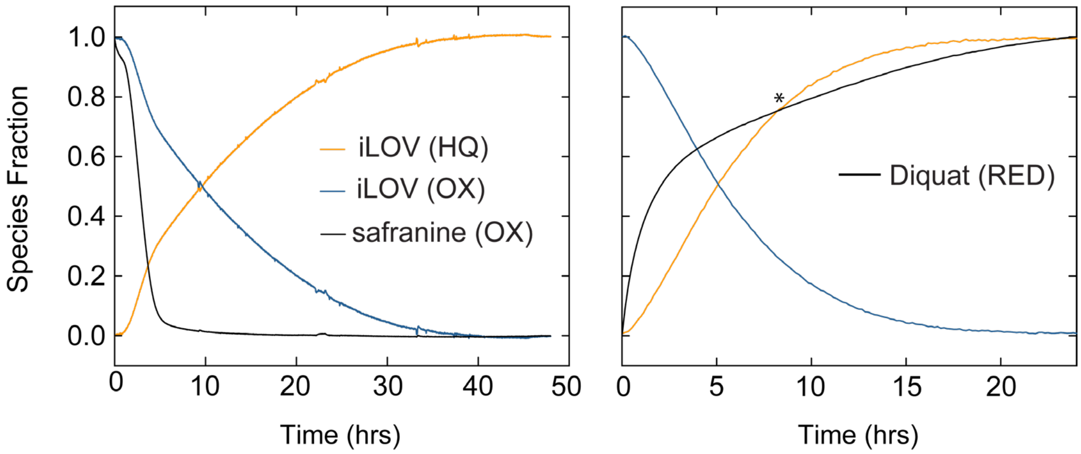


**Figure S9**. Xanthine/xanthine oxidase redox titration assay to measure the E°$_{OX/NHQ}$ couple. Reactions were performed in a VAC glovebox using the same spectrometer used for comproportionation experiments. For the reaction with safranine, the cuvette contained 169 µM xanthine, 0.3 U/L (10.2 µg/mL) xanthine oxidase, 2 µM $MV^{2+}$ as a mediator, 10 µM safranine, and 20 µM iLOV ($FMN_{OX}$) in 50 mM $P_i$ and 200 mM NaCl at pH 7.9. The fraction of oxidized iLOV was monitored at 420 nm, while the fraction of oxidized safranine was monitored at 520 nm. The fraction of reduced iLOV was assumed to be 1 – $FMN_{OX}$. The reduction of the standard dye proceeded far faster than the equilibration between the species. For the reaction with diquat as a standard, 130 µM of diquat, 30 µM of protein, 400 µM of xanthine, and 0.23 U/L (7.6 µg/mL) of xanthine oxidase was used in the same buffer adjusted to pH 7.9. The concentration of iLOV (OX) was monitored at 475 nm to minimize convolution with diquat, while the concentration of diquat (RED) was monitored at 777 nm. As before, the fraction of iLOV (NHQ) was assumed to equal 1 – $FMN_{OX}$. In this reaction, the fraction of reduced diquat starts higher than that of iLOV, but at some point in the titration the concentration of reduced iLOV becomes the major species. The reduction of diquat in this assay had an unexpected bi-phasic shape. Together these indicate that the equilibration between the two species was not at equilibrium, and that the oxidation of diquat while initially fast, became retarded as it became consumed by the slower iLOV (OX).

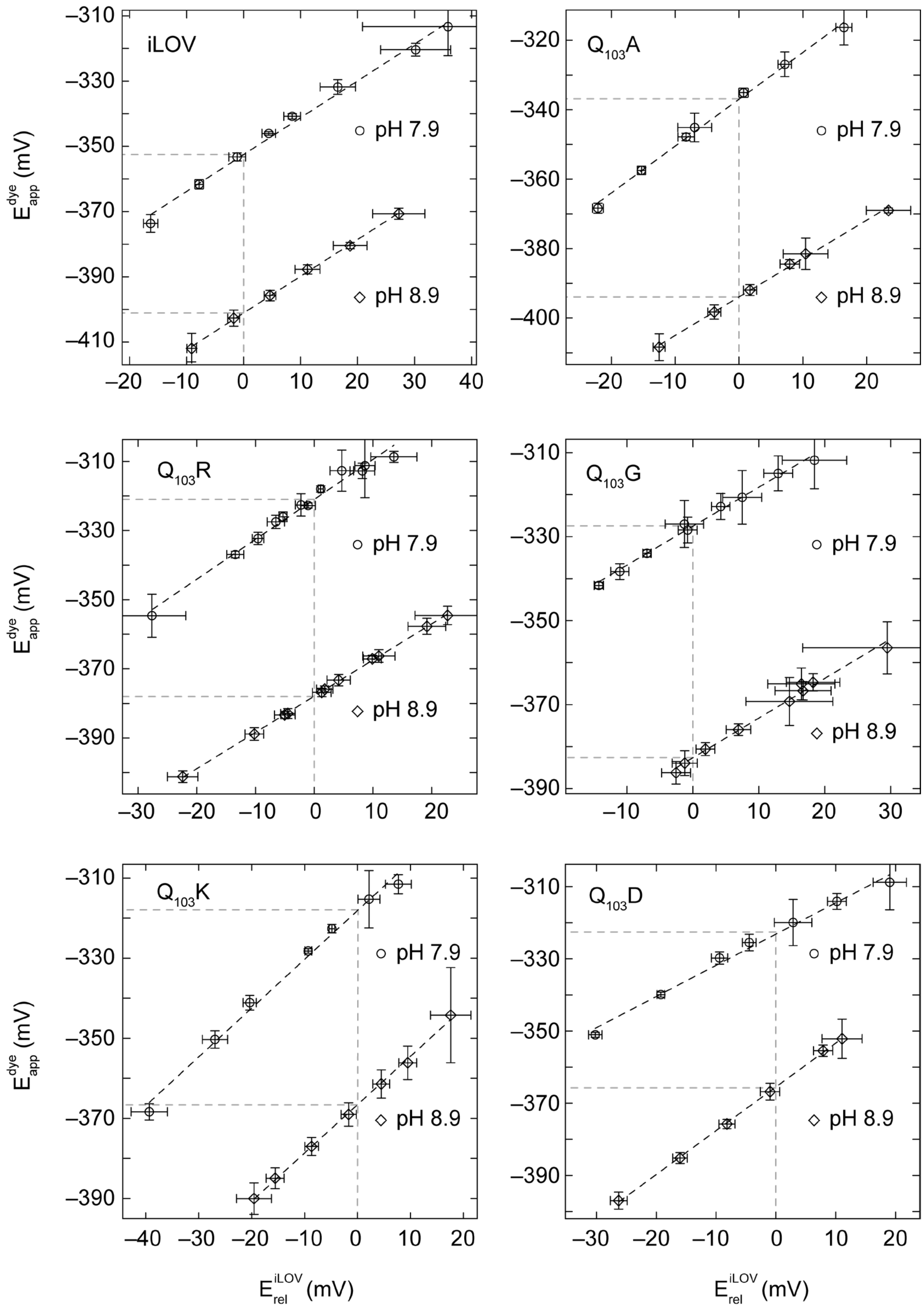


**Figure S10**. Redox titrations of iLOV and $Q_{103}X$ mutants. Data for iLOV is reproduced from the main text Figure 3. Titrations were conducted at pH 7.9 (circles) and 8.9 (diamonds). The experimental data were fit to a linear function (black dashed lines), and apparent E°OX/NHQ potentials are indicated on the y-axis (gray dashed lines). The results are summarized in the main text Table 1. Each data point represents the average of triplicate measurements in separate wells for the same conditions.

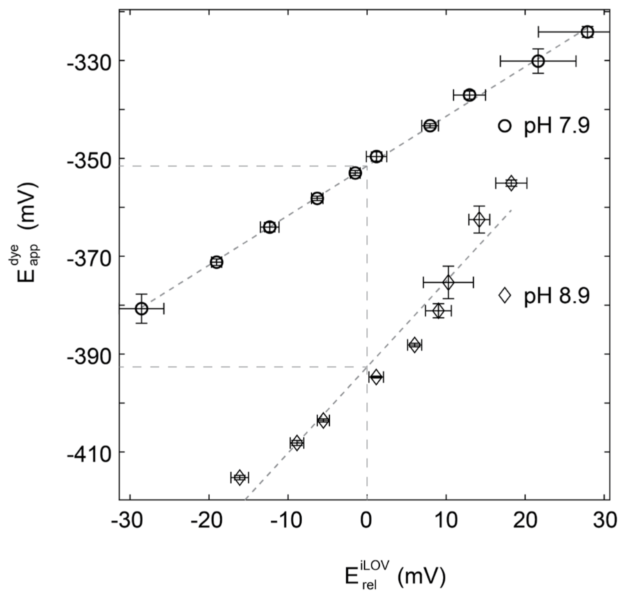


**Figure S11**. Oxidative redox titrations of iLOV with ferricyanide. The results of this titration recreate those of the reductive titration shown in Figure S9 and Main Text Figure 3. Significantly more scatter is observed in the pH 8.9 data, but the observed reduction potentials remain similar.

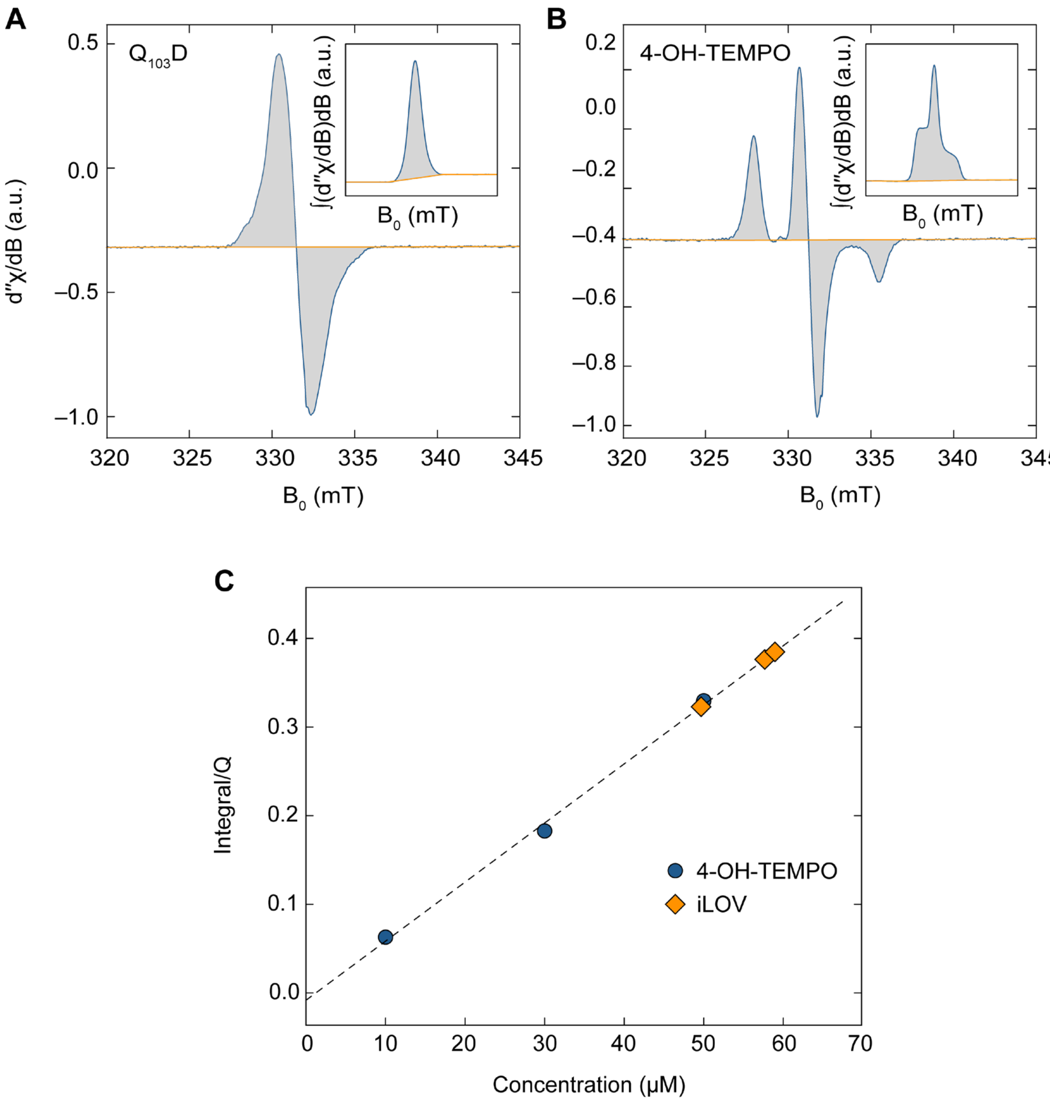


**Figure S12**. Radical quantification by EPR. **A** Representative EPR spectrum of photochemically reduced $Q_{103}D$ (gray shaded blue line). A linear baseline (orange) was subtracted from the spectrum, then integrated to yield the integrated spectrum in the inset. A second baseline was subtracted to account for the spectral asymmetry of the original spectrum (orange) and integrated again to yield the apparent total radical content. **B** Representative EPR spectrum of 4-hydroxy-TEMPO, treated identically to $Q_{103}D$. The 4-hydroxy-TEMPO standard concentration was determined by UV-vis to construct the calibration curve in C (blue circles), and compared to $Q_{103}D$ integrations (orange diamonds).

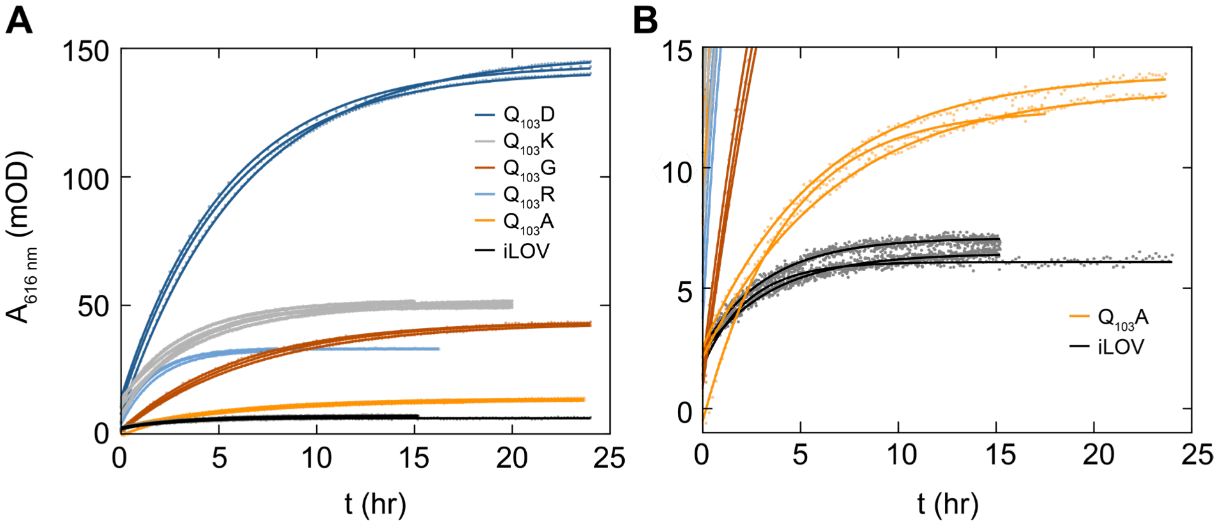


**Figure S13.** UV-vis quantitation of NSQ formation during the time course of comproportionation experiments for iLOV and Q103X mutants. **A** Absorbance at 616 nm ($A_{616\ nm}$, NSQ) data collected for each variant (colored dots) until equilibrium was reached, and single exponential fit (solid colored lines) to the approach to equilibrium. Experiments were performed and fit in triplicate to determine standard deviations. Equilibrium NSQ concentrations were determeined from the asymptote of the exponential fit. **B** Expanded view of iLOV and Q103A for clarity.

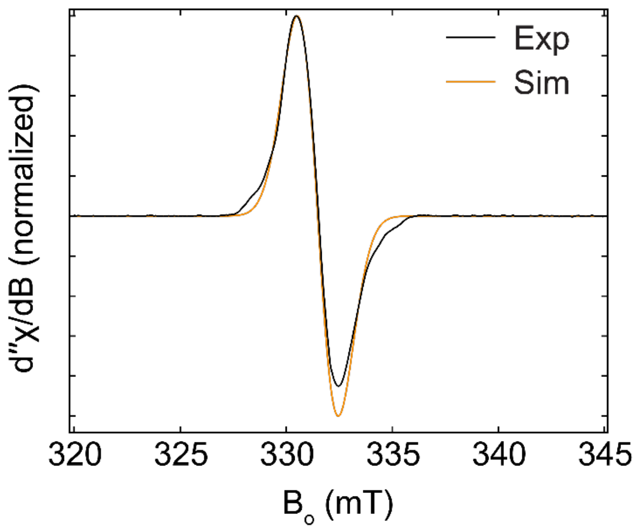


**Figure S14**. EPR spectrum of $Q_{103}D$ at pH 9.25. No significant differences in this radical spectrum are observed when compared to $Q_{103}D$ at pH 7.9. It is likely that the $pK_a$ of the N5 position in NSQ is >9.

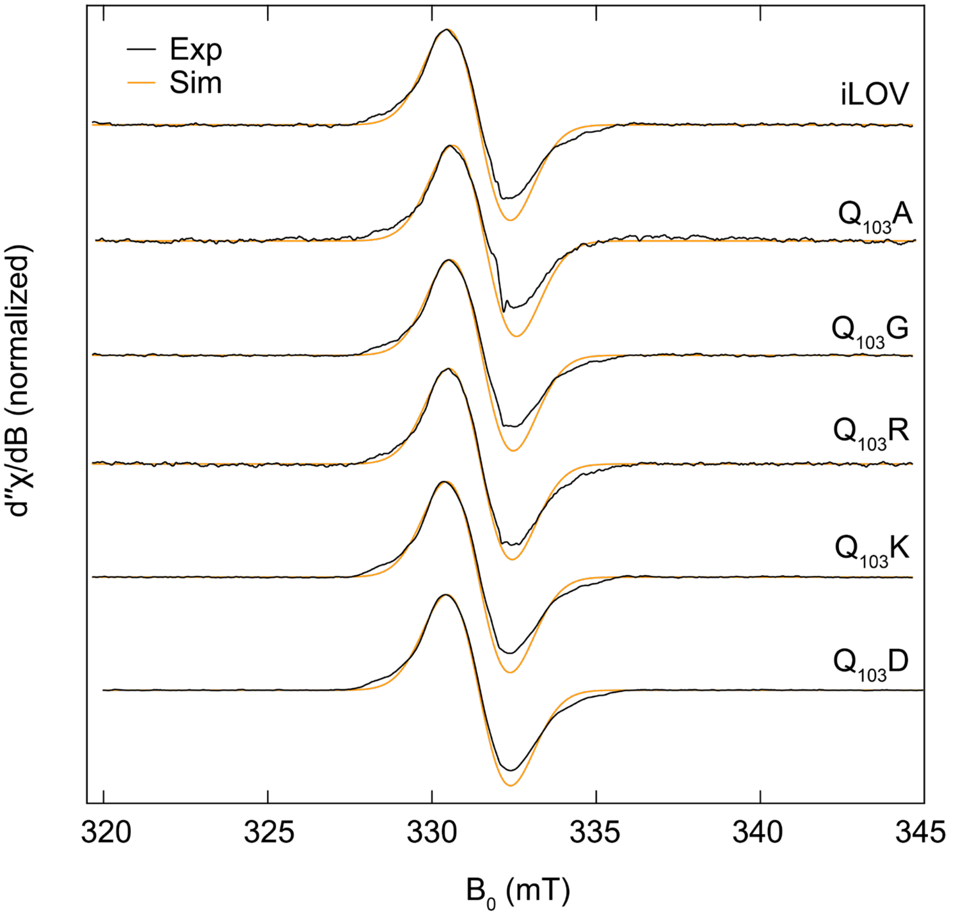


**Figure S15**. Continuous wave X-band EPR spectra of the NSQ radical of each iLOV mutant. Extracted simulation parameters are depicted in Table S5. Each spectrum remained identical.

**Table S5.** EPR simulation parameters for iLOV and $Q_{103}X$ NSQ spectra, and associated fit 68% confidence interval.

| Variant | $g_{iso}$ | Line width (mT) |
|---|---|---|
| iLOV | 2.0041 (1) | 1.92 (1) |
| $Q_{103}A$ | 2.0043 (1) | 1.94 (2) |
| $Q_{103}G$ | 2.0042 (1) | 1.92 (1) |
| $Q_{103}R$ | 2.0043 (1) | 1.93 (1) |
| $Q_{103}K$ | 2.0042 (1) | 1.94 (1) |
| $Q_{103}D$ | 2.0042 (1) | 1.94 (1) |
| $Q_{103}D$ (pH 9.5) | 2.0043 (1) | 1.94 (1) |

**Table S6**. ΔE values for each mutant using both the UV-Vis estimated extinction coefficient of 8,200 $M^{-1}$ $cm^{-1}$ and using the extinction coefficient estimated using EPR radical counting of 12,400 $M^{-1}$ $cm^{-1}$ for the NSQ radical. Standard deviations are shown in parentheses. While shifts of approximately 25 mV can be observed, the overall trends remain the same.

| iLOV Variant | UV-vis ΔE (mV) | EPR ΔE (mV) |
|---|---|---|
| WT | –212 (4) | –236 (4) |
| $Q_{103}A$ | –176 (3) | –195 (3) |
| $Q_{103}G$ | –112 (1) | –133 (1) |
| $Q_{103}R$ | –127 (1) | –149 (1) |
| $Q_{103}K$ | –104 (1) | –126 (1) |
| $Q_{103}D$ | –44 (1) | –68 (1) |

**Table S7.** Free energies of mutation computed for the oxidized (OX) and neutral semiquinone (NSQ) states of iLOV from different replicas. Indicated in the table are the pre-equilibration time (for λ=0 step) and the proximity of the water solvent to the N5 atom of flavin, measured as the start of the rise of the radial distribution function (RDF). Free energies highlighted in yellow indicate a "water out" value. Free energies highlighted in red are excluded due to a too-short pre-equilibration for that mutation.

| Mutant | State | Rep # | Eq. time (ns) | RDF rise (Å) | ΔG (kcal/mol) |
|---|---|---|---|---|---|
| **$Q_{103}D^{(0)}$** | OX | 1 | 50 | 1.9 | 22.9 |
| | OX | 2 | 50 | 4.6 | 22.0 |
| | OX | 3 | 3 | 4.5 | 22.4 |
| | NSQ | 1 | 3 | 2.7 | 21.5 |
| | NSQ | 2 | 3 | 2.8 | 20.0 |
| | NSQ | 3 | 50 | >5 | 20.8 |
| | OX | 1 | 3 | 2.3 | -77.2 |
| **$Q_{103}D^{(-)}$** | OX | 2 | 3 | 2.7 | -74.8 |
| | OX | 3 | 3 | 1.9 | -80.8 |
| | OX | 4 | 50 | 4 | -74.7 |
| | NSQ | 1 | 3 | 2.5 | -75.4 |
| | NSQ | 2 | 3 | 2.8 | -74.7 |
| | OX | 1 | 3 | 2.2 | 71.9 |
| **$Q_{103}A$** | OX | 2 | 3 | 4.8 | 72.1 |
| | OX | 3 | 50 | 4.8 | 73.2 |
| | NSQ | 1 | 3 | >5 | 69.3 |
| | NSQ | 2 | 3 | 2.6 | 71.9 |
| | NSQ | 3 | 50 | 2.5 | 72.3 |
| | OX | 1 | 3 | 1.8 | 61.1 |
| **$Q_{103}G$** | OX | 2 | 50 | 4 | 62.0 |
| | OX | 3 | 50 | 2.65 | 58.0 |
| | NSQ | 1 | 3 | 2.69 | 61.1 |
| | NSQ | 2 | 3 | 2.68 | 59.4 |
| | NSQ | 3 | 50 | 2.66 | 62.2 |
| | OX | 1 | 3 | 2.8 | -19.3 |
| **$Q_{103}K^{(+)}$** | OX | 2 | 50 | 2.4 | -11.5 |
| | OX | 3 | 50 | >5 | -15.3 |
| | OX | 4 | 100 | 2 | -11.9 |
| | NSQ | 1 | 3 | 2.6 | -8.2 |
| | NSQ | 2 | 3 | 2.6 | -12.8 |
| | NSQ | 3 | 50 | 2.68 | -16.3 |
| | NSQ | 4 | 100 | 2.7 | -14.9 |
| | OX | 1 | 3 | 4.1 | -170.3 |
| **$Q_{103}R^{(+)}$** | OX | 2 | 3 | 1.7 | -176.2 |
| | OX | 3 | 50 | >5 | -174.9 |

| | | | | |
|---|---|---|---|---|
| OX | 4 | 50 | 1.9 | -174.7 |
| OX | 5 | 100 | 1.87 | -172.6 |
| NSQ | 1 | 3 | 2.7 | -173.9 |
| NSQ | 2 | 3 | >5 | -174.0 |
| NSQ | 3 | 50 | 2.8 | -175.6 |

**Table S8:** Reduction potentials computed for $Q_{103}X$ mutations using different binning for the replicas obtained in **Table S7**. "All" uses all replicas. For $Q_{103}K^{(+1)}$ and $Q_{103}R^{(+1)}$, an extra bin after removing short (3 ns) pre-equilibration trajectories was included. Category 1 includes "water-in" models for both the OX and NSQ state. Category 2 includes "water-out" models for both the OX and NSQ states. Category 3 includes "water-out" for the OX state but "water-in" for the NSQ state. The average reduction potential and standard deviations are reported for different replicas belonging to each bin, as well as the error relative to the experimental reference. The bin in green displays the data reported in Table 3 of the manuscript.

| $Q_{103}D^{(0)}$ | ⟨Q⟩ (kcal/mol) | σQ (kcal/mol) | ⟨NSQ⟩ (kcal/mol) | σNSQ (kcal/mol) | $E_{red}$ (mV) | Total σ (mV) | Error (mV) |
|---|---|---|---|---|---|---|---|
| All | 22.4 | 0.5 | 20.8 | 0.7 | -386 | 52 | -44 |
| Category 1 | 22.9 | N/A | 20.7 | 1.0 | -364 | 45 | -22 |
| Category 2 | 22.2 | 0.3 | 20.8 | N/A | -399 | N/A | -57 |
| Category 3 | 22.2 | 0.3 | 20.7 | 1.0 | -395 | 59 | -53 |
| Exp. | | | | | -342 | | |

| $Q_{103}D^{(-)}$ | ⟨Q⟩ (kcal/mol) | σQ (kcal/mol) | ⟨NSQ⟩ (kcal/mol) | σNSQ (kcal/mol) | $E_{red}$ (mV) | Total σ (mV) | Error (mV) |
|---|---|---|---|---|---|---|---|
| All | -76.8 | 2.9 | -75.0 | 0.5 | -537 | 147 | -195 |
| Category 1 | -77.6 | 3.0 | -75.0 | 0.5 | -569 | 154 | -227 |
| Category 2 | -74.7 | N/A | N/A | N/A | N/A | N/A | N/A |
| Category 3 | -74.7 | N/A | -75.0 | 0.5 | -442 | 23 | -100 |
| Exp. | | | | | -342 | | |

| $Q_{103}A$ | ⟨Q⟩ (kcal/mol) | σQ (kcal/mol) | ⟨NSQ⟩ (kcal/mol) | σNSQ (kcal/mol) | Ered (mV) | Total σ (mV) | Error (mV) |
|---|---|---|---|---|---|---|---|
| All | 72.4 | 0.7 | 71.2 | 1.6 | -404 | 101 | 23 |
| Category 1 | 71.9 | N/A | 72.1 | 0.3 | -467 | 13 | -40 |
| Category 2 | 72.7 | 0.8 | 69.3 | N/A | -313 | N/A | 114 |
| Category 3 | 72.7 | 0.8 | 72.1 | 0.3 | -433 | 46 | -6 |
| Exp. | | | | | -427 | | |

| $Q_{103}G$ | ⟨Q⟩ (kcal/mol) | σQ (kcal/mol) | ⟨NSQ⟩ (kcal/mol) | σNSQ (kcal/mol) | $E_{red}$ (mV) | Total σ (mV) | Error (mV) |
|---|---|---|---|---|---|---|---|

| | | | | | | | |
|---|---|---|---|---|---|---|---|
| All | 60.4 | 2.1 | 60.9 | 1.4 | -482 | 155 | -101 |
| Category 1 | 59.5 | 2.2 | 60.9 | 1.4 | -518 | 160 | -137 |
| Category 2 | 62.0 | N/A | N/A | N/A | N/A | N/A | N/A |
| Category 3 | 62.0 | N/A | 60.9 | 1.4 | -410 | 63 | -29 |
| Exp. | | | | | -381 | | |

| **$Q_{103}K^{(+)}$** | **⟨Q⟩ (kcal/mol)** | **σQ (kcal/mol)** | **⟨NSQ⟩ (kcal/mol)** | **σNSQ (kcal/mol)** | **$E_{red}$ (mV)** | **Total σ (mV)** | **Error (mV)** |
|---|---|---|---|---|---|---|---|
| All | -14.5 | 3.7 | -13.0 | 3.5 | -520 | 312 | -76 |
| All, no 3ns | -12.9 | 2.1 | -15.6 | 1.0 | -340 | 133 | 32 |
| Category 1 | -11.7 | 0.3 | -15.6 | 1.0 | -288 | 55 | 84 |
| Category 2 | -15.3 | N/A | N/A | N/A | N/A | N/A | N/A |
| Category 3 | -15.3 | N/A | -15.6 | 1.0 | -444 | 42 | -72 |
| Experiment | | | | | -372 | | |

| **$Q_{103}R^{(+)}$** | **⟨Q⟩ (kcal/mol)** | **σQ (kcal/mol)** | **⟨NSQ⟩ (kcal/mol)** | **σNSQ (kcal/mol)** | **$E_{red}$ (mV)** | **Total σ (mV)** | **Error (mV)** |
|---|---|---|---|---|---|---|---|
| All | -173.7 | 2.3 | -174.5 | 1.0 | -426 | 143 | 0 |
| All (1&3), no 3ns | -174.1 | 1.3 | -175.6 | N/A | -391 | 56 | -5 |
| Category 1 | -173.6 | 1.5 | -175.6 | N/A | -373 | 66 | 13 |
| Category 2 | -174.9 | N/A | -174.0 | N/A | -498 | N/A | -112 |
| Category 3 | -174.9 | N/A | -175.6 | N/A | -427 | N/A | -41 |
| Exp. | | | | | -386 | | |

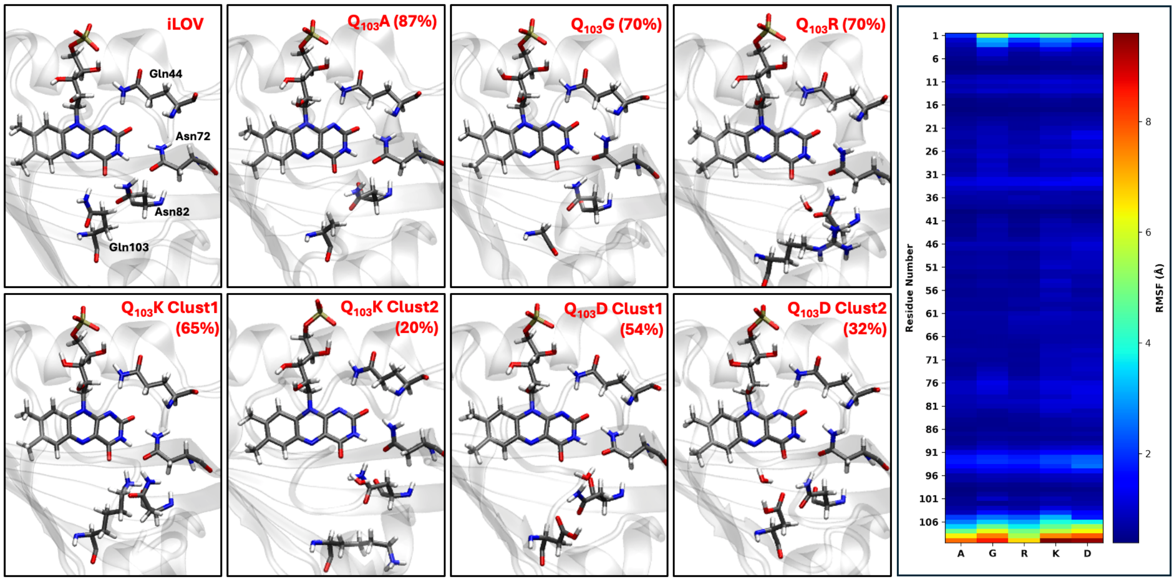


**Figure S16.** Left 6 panels: Structures obtained by clustering analysis for the mutants shown. The most populated cluster model is shown in each case, except for $Q_{103}D^{(0)}$ where we instead present the second most populated cluster model (the most populated one has 54% contribution with the aspartic acid pointing away from the flavin, but still has a water inside the binding pocket). The percentages indicate the number of frames belonging to the corresponding cluster model. Right panel: Heat map for the individual residue RMSF for the OX state of different mutants.
